# A murine aging cell atlas reveals cell identity and tissue-specific trajectories of aging

**DOI:** 10.1101/657726

**Authors:** Jacob C. Kimmel, Lolita Penland, Nimrod D. Rubinstein, David G. Hendrickson, David R. Kelley, Adam Z. Rosenthal

**Author notes:** Equal Contribution (Lolita Penland, Nimrod D. Rubinstein). DuPont Nutrition and Biosciences, 200 Powder Mill Rd, Wilmington, DE, 19803.

## Abstract

**Background:** Aging is a pleiotropic process affecting many aspects of organismal and cellular physiology. Mammalian organisms are composed of a constellation of distinct cell type and state identities residing within different tissue environments. Due to technological limitations, the study of aging has traditionally focused on changes within individual cell types, or the aggregate changes across cell types within a tissue. The influence of cell identity and tissue environment on the trajectory of aging therefore remains unclear.

**Results:** Here, we perform single cell RNA-seq on >50,000 individual cells across three tissues in young and aged mice. These molecular profiles allow for comparison of aging phenotypes across cell types and tissue environments. We find transcriptional features of aging common across many cell types, as well as features of aging unique to each type. Leveraging matrix factorization and optimal transport methods, we compute a trajectory and magnitude of aging for each cell type. We find that cell type exerts a larger influence on these measures than tissue environment.

**Conclusion:** In this study, we use single cell RNA-seq to dissect the influence of cell identity and tissue environment on the aging process. Single cell analysis reveals that cell identities age in unique ways, with some common features of aging shared across identities. We find that both cell identities and tissue environments exert influence on the trajectory and magnitude of aging, with cell identity influence predominating. These results suggest that aging manifests with unique directionality and magnitude across the diverse cell identities in mammals.

## Introduction

Aging is a gradual process of functional and homeostatic decline in living systems. This decline results in increased mortality risk and disease prevalence, eventually resulting in death. Aging appears to be a conserved feature of eukaryotic biology, affecting organisms as phylogenetically diverse as the single celled *S. cerevisiea*, the eutelic nematode *C. elegans*, mice, and humans [45, 63, 91]. Despite the near universal nature of the aging process, the underlying causes of aging are poorly understood. Aging phenotypes have been observed and hypotheses have been proposed for more than a hundred years [97, 50, 37, 64], but we do not yet know the cellular and molecular players that cause aging or how they differ between biological contexts. Both the fundamental nature of aging and its negative effects provide motivation to enumerate these players and establish causal relationships among aging phenotypes.

Mammalian aging phenotypes manifest at the organismal, tissue, cellular, and molecular levels [100]. Extensive research has produced catalogs of aging phenotypes at the physiological level, providing functional and behavioral hallmarks of age related decline. Likewise, molecular profiling of nucleic acids, proteins, and metabolites has provided a phenotypic description of aging in individual tissues [88, 42, 4, 56, 12].

Additional lines of inquiry have worked to address a classical question of aging biology – do different tissues age in the same way? Transcriptomic analysis at the bulk tissue level have revealed common traits of aging, as well as tissue-specific features [82, 43, 12]. Proteomic analysis of brain and liver in young and old mice similarly suggests that most age related changes are tissue-specific [72]. However, the cellular origins of aging phenotypes within a tissue remain largely unknown [12, 70, 78].

Our current understanding of aging phenotypes at the cellular level is less complete than at the tissue level. Mammals contain a multitude of distinct cell identities, each exhibiting specialized functions. In the mouse alone, recent cell atlas efforts have revealed more than 100 cell types [80, 39]. These surveys have catalogued diverse murine cell identities, but the plasticity of these identities and their contributions to tissue and organism level pathology remain unknown.

At both the molecular and functional level, a host of aging phenotypes and associated mechanisms have been revealed in individual cell types [23, 18, 35, 46, 48, 84, 20, 62]. While some of these studies present unique features of aging within individual cell identities, it is difficult to compare them systematically due to differences in experimental conditions and assay methodology. Using traditional molecular biology assays, it is difficult to measure high-dimensional molecular phenotypes across multiple cell identities, making large scale comparisons of aging phenotypes across cell identities intractable. The recent development of single cell RNA-sequencing (scRNA-seq) has ameliorated this limitation, allowing for measurement of transcriptional features across all prevalent cell identities in a tissue in a single experiment.

Although the technology has only recently matured, scRNA-seq experiments in individual tissues have already revealed novel aspects of the aging process. In a pioneering single cell RNA-sequencing study of hematopoietic progenitors, the axis of aging was shown to be opposite the axis of differentiation [53]. Multiple investigations have reported that cell-cell heterogeneity [34, 4] and gene expression variance [67] increase with age. However, the specific influence of cell identity and tissue environment on the trajectory and magnitude of aging has yet to be resolved.

Here, we employ scRNA-seq to generate a set of molecular profiles in which we can compare aging phenotypes across cell identities. By profiling 50,000+ cells from three tissues in young and old mice, we identify common features of aging that span cell identities, as well as features unique to each identity. Using matrix factorization and optimal transport methods, we compute trajectories of aging for each cell identity and assess the influence of identity and environment on these trajectories.

## Results

### Single cell RNA-sequencing identifies a diversity of cell types and states in young and old mouse tissue

We collected transcriptional profiles of young and old cells of many identities by isolating single cells from the kidney, lung, and spleen of *n* = 4 young (7 months) and *n* =3 old (22-23 months) *C57Bl/6* mice. All three tissues were collected from the same animals. Isolations were performed at the same time of day for each animal, limiting circadian variation which affects the expression of nearly half of all murine genes [101]. After single cell isolation, cells were immediately encapsulated and barcoded for library preparation using the 10X Genomics microfluidics system, followed by subsequent sequencing. Using standard techniques for the identification of cell containing microfluidic droplets, we recovered 55,293 individual cell transcriptomes (see Methods).

We determined cell type and state identity by leveraging annotations provided in the *Tabula Muris* compendium [80]. These annotations provide labels at the cell type level and follow the structured hierarchy of the “cell ontology” [8]. Some age-related changes may be unique to individual states within a cell type. Similarly, some changes at the level of cell types may be mediated by differences in cell state proportions (i.e. CD4 vs. CD8 T cells). To ensure that we can explicitly detect these cell state level changes, we manually annotated cell states within each cell type in the *Tabula Muris* (see Methods, Supp. Fig. 1). We use the term “cell identity” to refer to the combination of cell type and state labels, such that CD4 T cells and CD8 T cells are different cell identities (Fig. 1).

**Figure 1:**
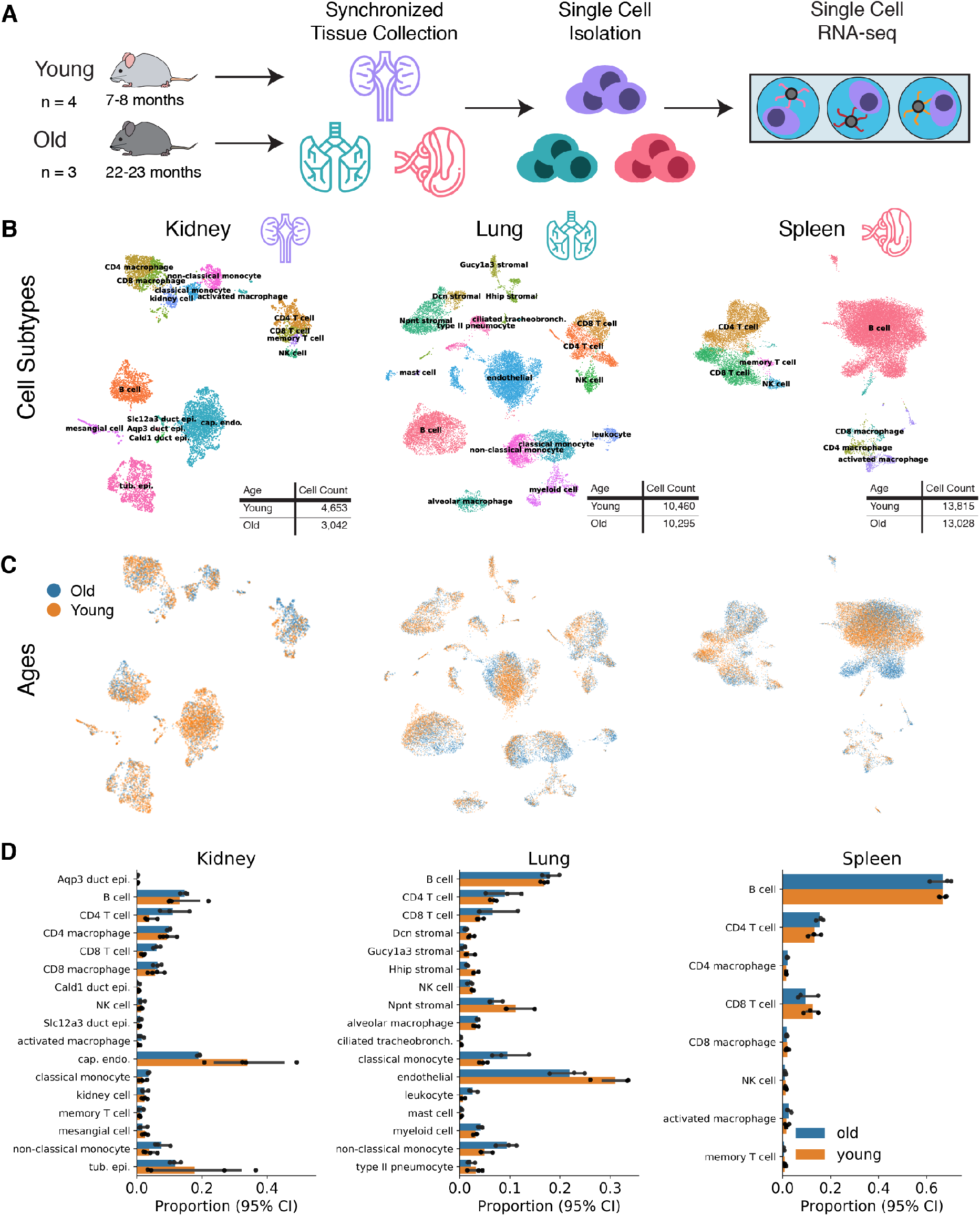
scRNA-seq reveals that non-immune cell type proportions are preserved with age. **(A)** Schematic representation of the experimental design. Kidney, lung, and spleen tissue were isolated simultaneously from each young and old mouse. After generating single cell suspensions, cells were prepared for scRNA-seq using the 10X Chromium system. **(B)** UMAP embeddings of each tissue investigated in our data set. Colors represent cell type annotations. Cell types are derived using a deep neural network classifier trained on the Tabula Muris data set. **(C)** Matching UMAP embeddings depicting the age of each cell. **(D)** For each animal, we computed the proportion of cells in each state. The mean proportion of each cell state for each age across animals is presented as a bar graph. The underlying proportions observed in individual animals are overlaid as black dots.

We trained deep neural networks to classify cell types based on these annotations, then used these networks to predict cell types in our data (Fig. 1A, B; see Methods). We found that the networks transfer labels with high fidelity. We validated classifications by inspecting marker gene expression *post-hoc* (Supp. Fig. 2, 3) and computing correlations between cell identities in our data and the *Tabula Muris* (Supp. Fig. 4). From these cell identity annotations, we identify 19 unique cell types and 28 unique cell states across the three tissues. Comparing our cell types to the *Tabula Muris*, we recover all but one of the cell types identified (kidney loop of Henle epithelial cells, Supp. Fig. 3, 5). The cell type proportions we recover differ from the *Tabula Muris* – e.g. we recover comparatively more immune cells in the kidney and lung – but are not outside expected range based on previous comparisons between single cell RNA-seq datasets [80, 74].

#### Immune cells are more prevalent in old kidneys and lungs, while non-immune cell type proportions are preserved

One prospective way in which aging may influence tissue function is by altering the proportion of each cellular identity within the tissue. This change in cell identity compositions may occur at either the level of cell types, or by shifting the distribution of cell states within a cell type. To investigate the former possibility, we quantified the proportion of each cell type within each tissue across ages.

In both the kidney and lung, lymphocytes were significantly more abundant in old animals compared to young animals (*t*-test on additive log-ratio transformed proportions, *q* < 0.05)(Fig. 1D). In old kidneys, we found a roughly 2 fold increase in T cells, classical monocytes, and non-classical monocytes, and in old lungs a corresponding 2 fold increase in classical and non-classical monocytes and a roughly 1.3 fold increase in T cells. This may reveal increasing immune infiltration of the the non-lymphoid tissues with age, as suggested in previous studies of kidney, lung, and other non-lymphoid tissues [70, 78, 92, 6, 65]. However, we can not rule out that our recovery of specific cell types may be confounded by an interaction of aging with our isolation procedures.

Considering only non-immune cells in the kidney and lung, cell type proportions were not substantially altered by aging (Supp. Fig. 6B). Likewise in the spleen, we found minimal change in the proportion of cell types between young and old animals (Fig. 1D). While changes in non-immune cell type proportions are subtle, we cannot rule out that even these subtle changes may influence the aging process.

Shifting cell state proportions within a cell type may be an alternative mechanism by which aging phenotypes manifest. Examples of this phenomenon are present in the literature, such as the observed decrease in naive CD8 T cells relative to other T cell states [29, 38] and the shift from highly regenerative to less regenerative stem cell states in the blood and muscle [23, 14]. To address whether cell state proportions change with age, we quantify the proportion of cells in a given cell state for each cell type.

Investigating spleen T cells, we recapitulate the finding that CD8 T cells are less abundant in old animals. We observed a shift in the population of kidney collecting duct epithelial cells – old animals exhibit a decreased frequency of *Cald1*+ collecting duct cells relative to *Slc12a3*+ collecting duct cells (*χ*^2^ contingency table, *q* < 0.05). *Aqp3*+ and *Slc12a3*+ cells are likely principle cells of the collecting duct based on expression of *Scnn1a*. *Cald1*+ cells are also marked by *Phgdh*, which has a reported mosaic expression pattern in the proximal tubule, and *Cryab*, which was similarly identified to mark a distinct subpopulation in the Mouse Microwell Atlas [39] (Supp. Fig. 6C). However, we are not aware of a defined role for this cell population, making it difficult to speculate on the impact of this shift in cell state proportions. Other cell types with notable cell state substructure do not show shifts with age, such as lung stromal cells (Supp. Fig. 6A).

#### Cycling cells are similarly rare in young and old animals

Previous reports have suggested that cell cycle activity changes with age in multiple cell populations. In blood progenitors, cell cycle kinetics are accelerated with age [53], while in muscle progenitors the frequency of cycling cells increases with age [23], and in the intestinal crypt cycling cells become less frequent [69]. To investigate the possibility of changes in cell cycle frequency in our data, we evaluated cell cycle activity by scoring the expression of S-phase associated genes and G_2_M-associated genes [90] (see Methods). We observe only subtle changes in either of these cell cycle module scores with age across cell identities. The proportion of putatively cycling cells is also very small across cell identities (Supp. Fig. 7). These results suggest that the cell cycle rate in cell identities we observe is not dramatically changed with age. However, we cannot discount the possibility that the cell cycle scoring method we use is insufficient to detect differences.

The accumulation of non-cycling senescent cells in aging tissues has been reported in several previous studies [25]. The reported magnitude of senescent cell accumulation varies between tissues. In aging mouse kidney, the proportion of cells with senescence associated *β*-galactosidase activity increased from roughly 0.2% to 1.2%, whereas in epicardial cells the proportion increases from 2% to 10% (12 months to 18 months) [7]. Similar observations have been made using paired-end single cell RNA-seq in the human pancreas, where the proportion of cells expressing senescence marker gene *CDKN2A* increases from roughly 7% to 15+% between early- (21-22 years) and mid-adulthood (38-54 years) [34]. To investigate whether senescent cells are more prevalent in the old tissues we observe, we similarly measured expression of *Cdkn2a*. We find that the *Cdkn2a* locus (p16-Ink4a and p19-Arf) is not significantly upregulated with age in any of the cell identities we observe (Supp. Fig. 8). Due to the overlapping nature of the p16-Ink4a and p19-Arf reading frames, we note that we cannot distinguish transcripts from these two proteins using 3′-end RNA-seq alone [44]. We also scored the activity of a curated set of senescence-associated genes using the AUCell approach [1], but we do not find large differences in this score with age (Supp. Fig. 8).

### Changes in cell-cell variation with age depend on cell identity

Single cell analysis allows us to measure not only the mean expression of each gene, but also the variation within a cell population. Previous work has suggested that both gene expression variance and cell-cell heterogeneity increase with age [67, 34, 5]. These two types of variation differ in subtle but important ways. Gene expression variance quantifies the mean dispersion across genes in the transcriptome, such that each gene contributes equally. Because genes are equally weighted, changes in gene expression variance are unlikely to be driven by a small number of genes. Likewise, multivariate differences in gene expression between cells are not resolved due to the focus on mean dispersion values. Increased gene expression variance may reflect a global change in transcriptional noise, perhaps due to loss of regulatory control, as suggested in previous aging studies [93].

By contrast, cell-cell heterogeneity measures the average distance in transcriptional space between cells in a population. These distances capture multivariate differences in gene expression, capturing variation in cell state that manifests across genes. They also account for gene expression level, such that a small number of more highly expressed genes can drive changes in cell-cell heterogeneity. Increased levels of cell-cell heterogeneity may reflect a diversification of cellular states within a population (Fig. 2A).

**Figure 2:**
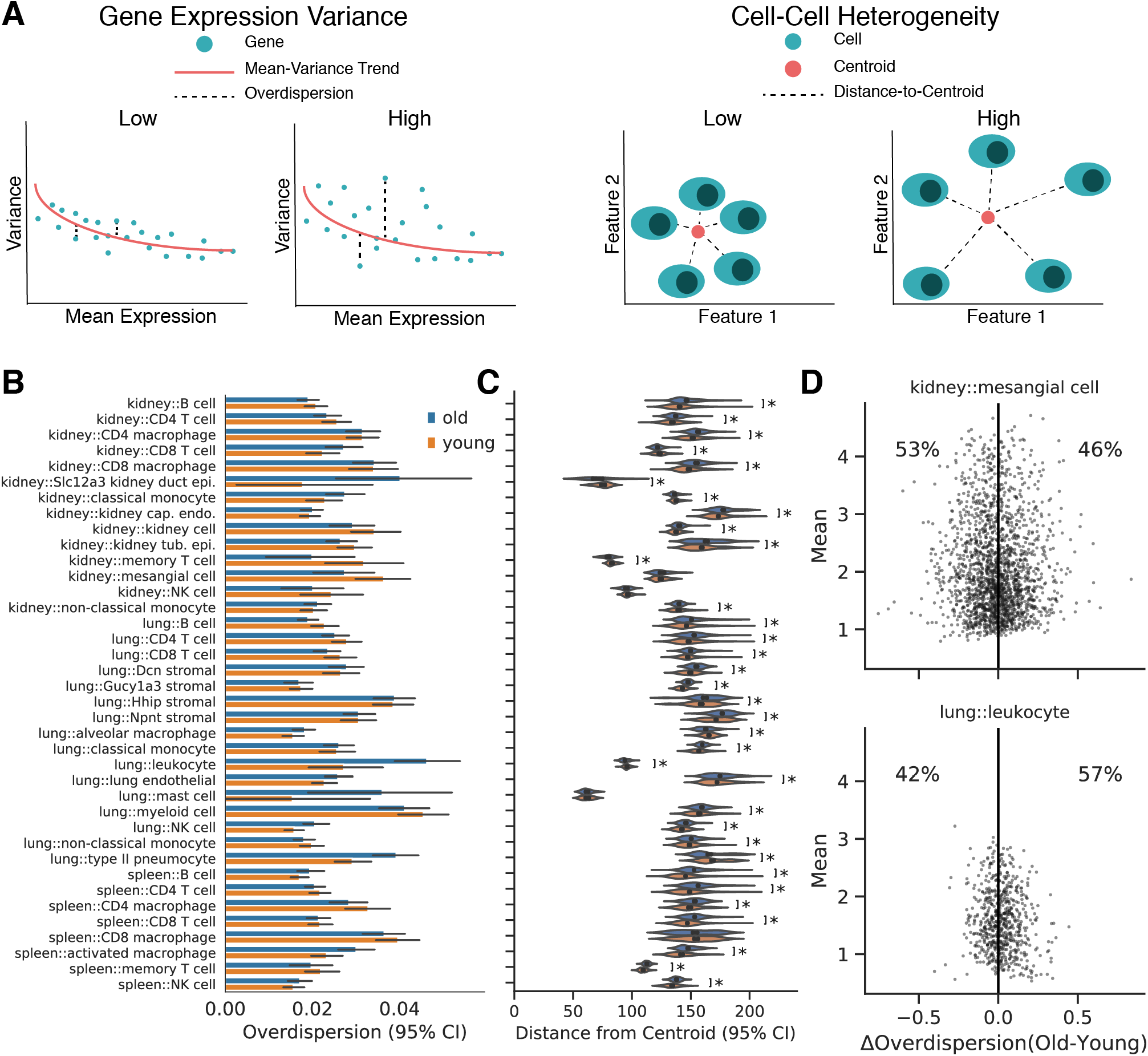
Changes in cell-cell heterogeneity with age depend on cell identity. **(A)** Diagram illustrating the difference between gene expression variance (left) and cell-cell heterogeneity metrics (right). Gene expression variance measures the variance of the average gene in a manner that controls for mean expression levels. Cell-cell heterogeneity measures the average difference between cells across genes. Examples of data providing high and low values for each metric are schematized. **(B)** Overdispersion values (computed using the difference from the median method) for each cell identity conditioned on age. Each point in the underlying data represents a single gene. Many cell identities do not show a substantial shift in the overdispersion distribution. Some identities show increases in the mean overdispersion with age, while others show decreases. **(C)** Cell-cell heterogeneity measurements based on Euclidean distances to the population centroid for each cell identity and age. Each point in the underlying data represents a single cell. Most cell identities exhibit increased cell-cell heterogeneity in old cells (*: Wilcoxon Rank Sums, *q* < 0.05). However, some identities show decreased heterogeneity with age as well (kidney::CD8 T cell, kidney::classical monocyte). **(D)** Difference in overdispersion between old and young cells (ΔOverdispersion) as a function of mean gene expression value in *Vim*+ kidney capillary endothelial cells (upper) and lung leukocytes (lower). Each point represents a single gene.

Both transcriptional variation and cell-cell heterogeneity have important implications for cell physiology and function of a cell population, as explored in seminal studies of transcriptional noise in cell fate selection [17, 89] and bet hedging [55, 2, 26, 85, 10]. To determine if age-related changes in variation and heterogeneity depend on cell identity and tissue environment, we evaluate both properties across the many combinations of cell identity and tissue environment we observe.

We evaluate transcriptional variation using the difference from the median (DM) method [51] to estimate “overdispersion.” Overdispersion refers to the residual variation in gene expression observed for a given gene after accounting for the mean:variance relationship in gene expression data [67] (see Methods). Across the cell identity/environment combinations we observe, we find that transcriptional variation by this metric is largely unchanged (Fig. 2B). Individual cell identities can be identified that exhibit both an increase in variation with age (lung leukocytes) or a decrease in variance with age (kidney mesangial cells) (Fig. 2D).

We quantified cell heterogeneity in each cell identity/environment combination using the distance to the centroid method as previously introduced [34, 5]. Cell-cell heterogeneity appears to increase for many cell identities (Wilcoxon Rank Sums, *q* < 0.05), including B cells across all three tissues, and lung stromal cells. We also observe decreased heterogeneity with age in some cell identities, such as lung type II pneumocytes and kidney CD8 T cells (Fig. 2C). Taken together, these results indicate that changes in gene expression variance and cell-cell heterogeneity with age are not universal and depend on cell identity. This suggests a more nuanced view of the notion that an increase in noise is a defining hallmark of aging at the single cell level [34, 93].

### Differential expression reveals aging phenotypes common across cell identities

Prior studies have revealed differentially expressed genes within whole tissues or individual cell types in aging [27, 78, 70, 12, 47, 98, 88, 43]. However, it remains unclear to what degree age-related transcriptional changes are shared or unique across cell identities. To address this outstanding question, we performed differential expression analysis within each cell identity between young and old mice.

We identified differentially expressed genes for each cell identity/tissue environment combination using the Rank Sums method (see Methods). For each differentially expressed gene, we counted how many cell identities differentially express that gene in the same direction. The majority of differentially expressed genes with age are specific to one or a few cell identities (Fig. 3A).

**Figure 3:**
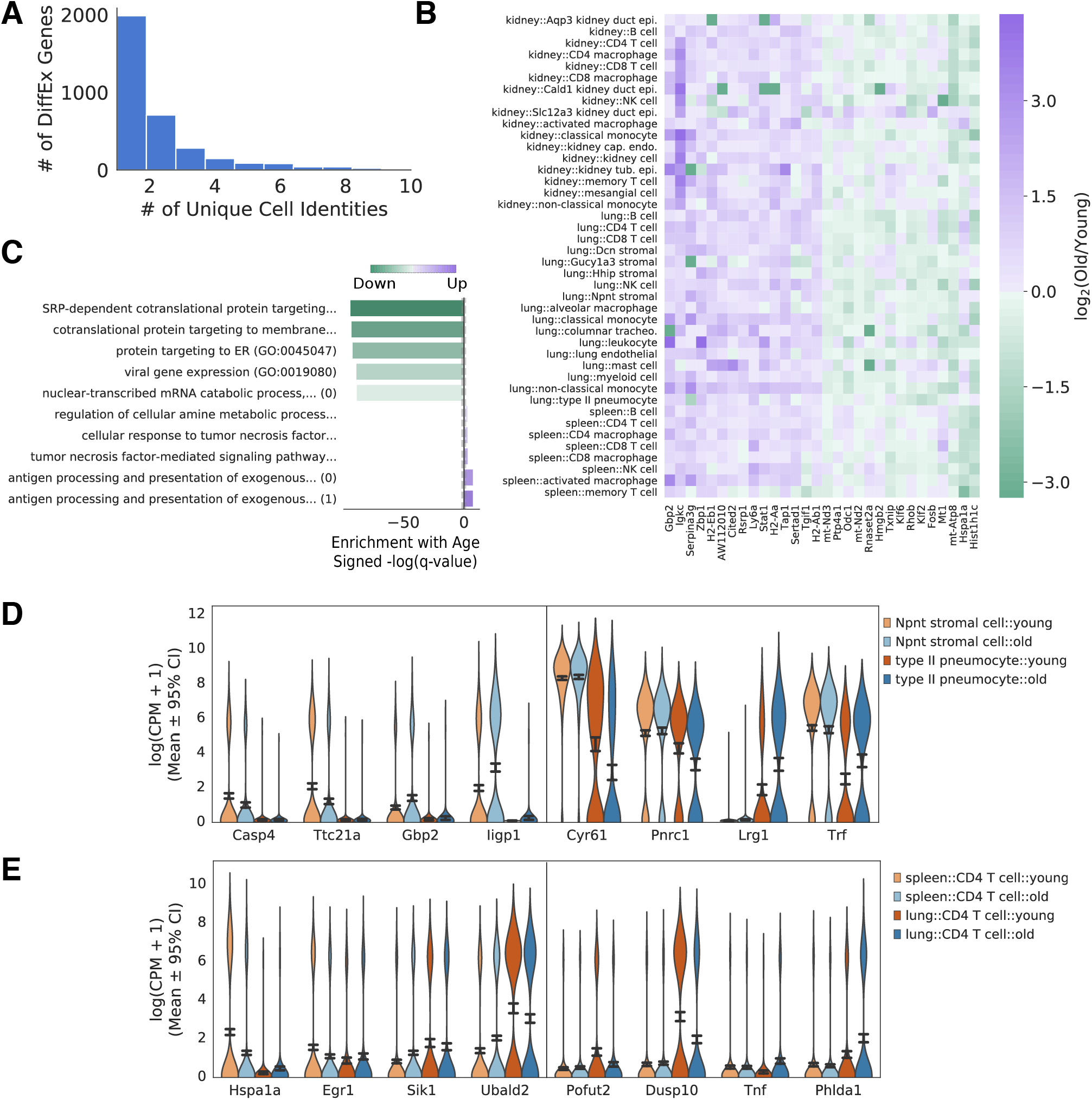
Differential expression analysis identifies common age related changes across cell identities and tissue environments. **(A)** The number of genes differentially expressed in at least *k* cell types, presented as a histogram. **(B)** Heatmap of top 15 genes significantly changed in the each direction across >= 5 cell types. Fold changes between old and young cells are presented for each cell type in each tissue. While no gene is universally changed across cell types, each gene is changed across multiple tissues and developmental lineages. **(C)** Top 5 enriched gene ontology terms for genes that are downregulated (negative values) and upregulated (positive values) with aging across >= 5 cell types. Dotted lines represent the *α* = 0.05 significance threshold. Antigen processing and metabolic pathways appear to be upregulated, while protein translation and translocation pathways appear to be downregulated with aging. **(D)** Violin plot of genes that are uniquely changed between two cell states in the lung (Wilcoxon Rank Sums, *q* < 0.05) – *Npnt* stromal cells and type II pneumocytes. Each gene presented is significantly upregulated or downregulated in one cell type and does not change in the same direction (log_2_ (Old/Young) < 0.1) in the other cell type. Confidence intervals are computed by bootstrapping. For each cell state, we show the top two specific downregulated genes and upregulated genes. **(E)** Violin plot of genes that are uniquely changed between CD4 T cells isolated from the spleen and the lung. Genes are selected as in **(D)**.

However, a subset of 261 genes are differentially expressed across > 5 cell identities and show consistent changes across multiple cell identity/tissue environment combinations (Fig. 3B). Here, we chose 5 cell identities as a cutoff for common differentially expressed genes to reduce the number of genes identified due to common differential expression in highly similar cell identities (i.e. monocytes and macrophages). Thus, some aspects of transcriptional aging are common to many cell identities. Using gene ontology enrichment analysis for biological processes [54], we identify SRP-dependent protein localization and protein translocation to the ER as commonly downregulated across cell identities. This observation is consistent with the observed interaction of protein translocation systems with aging [40, 87] (Fig. 3C, lower). Antigen processing and inflammatory pathways are significantly upregulated with age, a result that has likewise been observed across tissues [78, 70, 12, 72]. Performing hierarchical clustering on this common set of differentially expressed genes, we find gene clusters enriched for specific inflammatory processes (type I interferon signaling, cytokine secretion), suggesting that more than one immunological pathway is changed with age (Supp. Fig. 9).

Increases in inflammation associated gene transcription have previously been reported in multiple tissues. However, previous investigations have relied on bulk transcriptional assays, making it difficult to determine if inflammatory genes signatures were upregulated in all cells within a tissue, or if more immune cells had infiltrated the tissue [78, 70, 12]. To investigate the possibility that immune pathways are upregulated across many non-immune cell identities, we identified a set of genes that are differentially expressed in at least 3 non-immune cell identities. Gene ontology enrichment analysis on this gene set reveals that inflammatory gene sets (T cell activation, B cell activation, viral entry, response to cytokines) are upregulated, even in these non-immune cell identities. However, the enrichment is much less significant than when immune cells are included (Supp. Fig. 10). At the gene level, we find *Ikgc, Cd74*, and *B2m* are commonly upregulated with aging. *B2m* has previously been reported to increase in the aging systemic milieu, and reports have indicated it may play a causal role in aging brain pathology [86].

Many genes change expression uniquely in individual cell identities, even within the same tissue. For instance, *Npnt* lung stromal cells exhibit upregulated and downregulated genes that are unchanged in lung type II pneumocytes and vice-versa (Fig. 3D). Gene ontology enrichment analysis reveals that *Npnt* stromal cells downregulate fibrinolysis and muscle development gene sets, while upregulating responses to fluid shear stress that are unchanged in type II pneuomcytes. By contrast, type II pneumocytes downregulate degranulation associated gene sets that are unchanged in stromal cells (Supp. Fig. 10).

We also observe genes that are differentially expressed in a tissue-specific manner. For example, CD4 T cells in the lung demonstrate downregulated and upregulated genes that are unchanged in CD4 T cells of the spleen and vice-versa (Fig. 3E). Collectively, these results indicate that while most differentially expressed genes are specific to individual cell identities, a subset of changes appears to be common across many identities. This common subset includes previously reported increases in inflammatory gene expression, which we find in both immune and non-immune cells. Cell identity and tissue environment also influence the genes which are differentially expressed, as demonstrated in comparisons of different cell identities in the same tissue, or the same cell identity in different tissues.

### Aging manifests novel B cell states in the spleen

Beyond shifting cell state proportions, aging could promote the formation of novel cell states unseen in young animals. Age-associated cell states may then contribute to aging phenotypes at the cell type and tissue scales. We observe an example of a cell state that arises with age in spleen B cells.

Spleen B cells in our experiments exhibit three distinct clusters which do not map well to a canonical subtyping scheme (Fig. 4A). We observe that one of these clusters is dominated by cells from old animals (Fig. 4A, C; *p* < 0.05, *χ*^2^ contingency table). We performed differential expression analysis on each of these clusters to identify marker genes, and note high levels of *Apoe* in the larger cluster and high levels of *B2m* in the other. It has been reported that C57Bl/6 mice, like those in this study, die with an unusually high incidence of lymphoma [16, 21, 75]. To ask if these clusters dominated by old cells potentially represent lymphomas, we examined the expression of several genes associated with B cell lymphomas in a previous study [58]. We identify several lymphoma associated genes significantly upregulated in each of these clusters (Fig. 4B, rows 4-9)(*q* < 0.05, Wilcoxon Rank Sums test). Performing gene set enrichment analysis using the MSigDB Hallmark Gene Sets [61], we observe that *Myc* target genes and DNA repair pathways are upregulated, while p53 pathway genes are downregulated (Fig. 4D). This pattern of differential gene expression suggests that the *Apoe* high population is pre-neoplastic.

**Figure 4:**
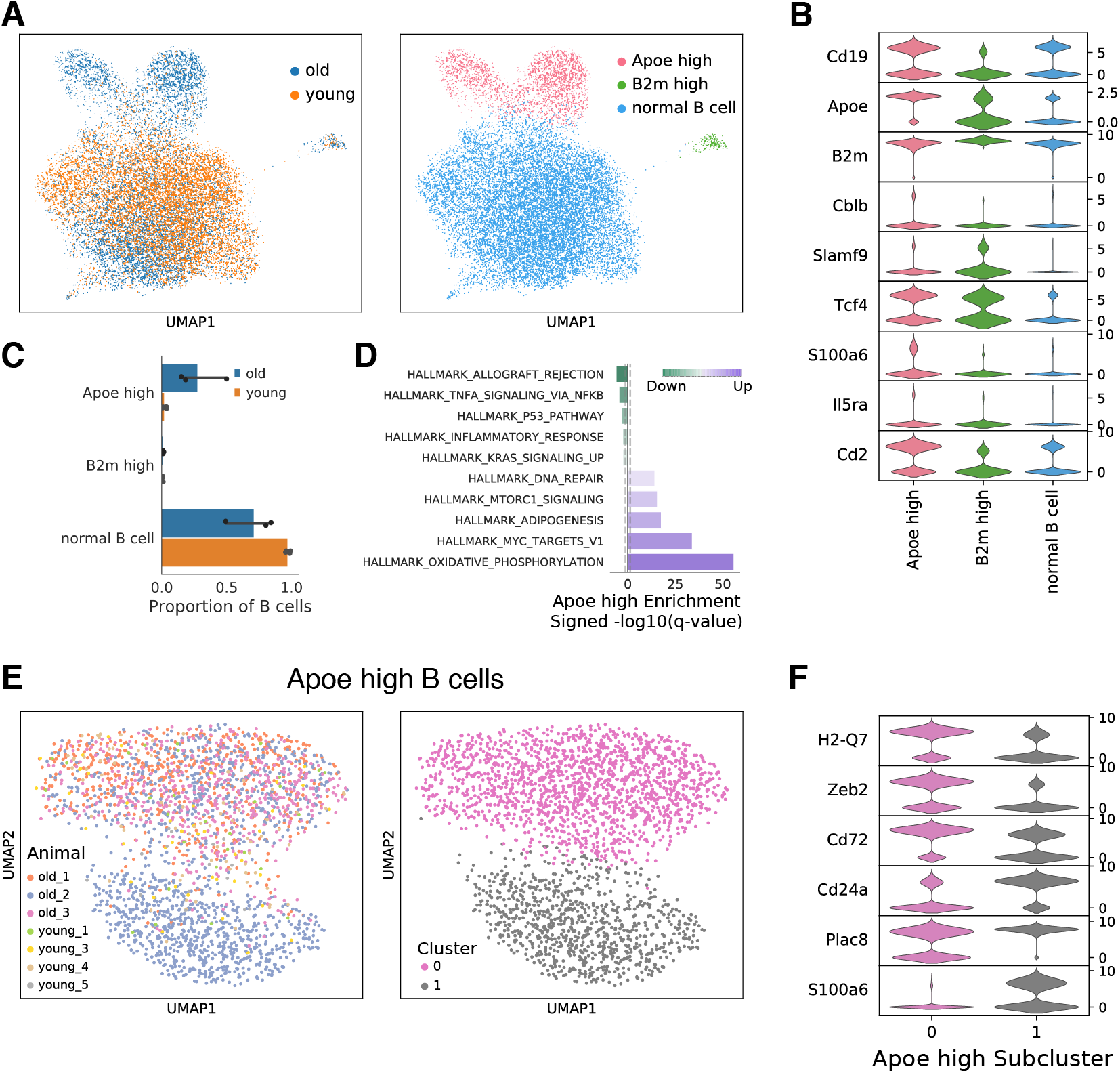
Aging manifests novel B cell states in the spleen. **(A)** UMAP embeddings of the spleen B cell compartment. Louvain clustering identifies three subpopulations within the B cell compartment of the spleen, named by their marker genes (left). The distribution of young and old cells is shown for comparison (right). **(B)** Marker genes for each cluster and a set of lymphoma specific genes presented as violin plots. All markers presented are significantly enriched in the *Apoe* high or *B2m* high cluster. **(C)** Proportions of cells in each cluster as a function of age. The *Apoe* high and *B2m* high states are occupied predominantly by aged cells (*t*-test on additive log ratio transformed proportions, *q* < 0.05). **(D)** Enrichment of MSigDB Hallmark gene sets based on differentially expressed genes in the *Apoe* high cluster. *Myc* targets, mTOR signaling, and DNA repair pathways are upregulated while p53 signaling is downregulated, suggestive of neoplasia. Gray dotted lines represent the *α* = 0.05 significance threshold. **(E)** UMAP embedding of the *Apoe* cell cluster. A second Louvain clustering iteration reveals two clusters within this *Apoe* high group (left). Visualizing the animal of origin for each cell, it is apparent that Cluster 1 is dominated by a single old animal (right). **(F)** Marker genes for each of the sub clusters within the *Apoe* high state presented as violin plots. Cluster 0 is enriched for lymphoma associated gene *Zeb2* while Cluster 1 is enriched for *Plac8*.

At a finer level of detail, we find that the *Apoe* high cluster appears to have two distinct lobes. One of these two lobes seems to contain cells almost entirely derived from a single old animal (1 of 3) (Fig. 4E), while the other lobe contains cells found in all of the old animals observed. To determine what differentiates these lobes, we examined marker genes and identify differences in the expression of *Zeb2*, the B cell marker *Cd72, Plac8*, and *Cd24a* as discriminating features (Fig. 4F; *q* < 0.05, Wilcoxon Rank Sums test). Each of these genes has previously been associated with neoplasia [102, 49, 60].

These emergent cell states support the notion that some aging phenotypes manifest not by shifting the distribution or location of youthful states, but by creating new states altogether. Neoplastic cells are an extreme case of such a phenomenon. The observation of a cell state that is generally ubiquitous in old mice and a related state that is present in only one animal also highlights the commonalities and the stochastic animal-animal heterogeneity that are associated with studies of the aging process.

### Cell identity determines the trajectory of aging

Our differential expression and heterogeneity analyses suggest that cell identities age differently, and that the same cell identity ages differently across tissues. How much do cell identity and tissue environment influence the trajectory of aging? To answer this question quantitatively, we compute a trajectory of aging for each cell identity in each tissue based on their transcriptional profiles.

We perform this analysis using an embedding space derived by non-negative matrix factorization (NMF). NMF embeddings of transcriptional data have been shown to recover relationships between genes, such that each component of the embedding may be interpreted as a gene expression program [52, 22, 83]. Likewise, values of each component in each cell can be interpreted as the activity of that program. By using an NMF embedding to compute trajectories of aging, we are able to make qualitative interpretations at a level of abstraction above individual genes.

To compute aging trajectories, we first embedded all cells observed across tissues in a 20-dimensional NMF space (Fig. 5B; Supp. Fig. 11; see Methods, dimensionality chosen based on the trade-off between interpretability and explained variance). To assign semantic meaning to the embedding dimensions, we identified genes associated with each dimension by thresholding on the dimension loadings and analyzed gene set enrichment (Supp. Fig. 12). We computed the aging trajectory for each cell identity/tissue combination as the distance between the centroid of the young cells and the centroid of the old cells in this embedding (see Methods). This procedure yields a 20-dimensional vector representing the trajectory of age-related change observed in each cell identity.

**Figure 5:**
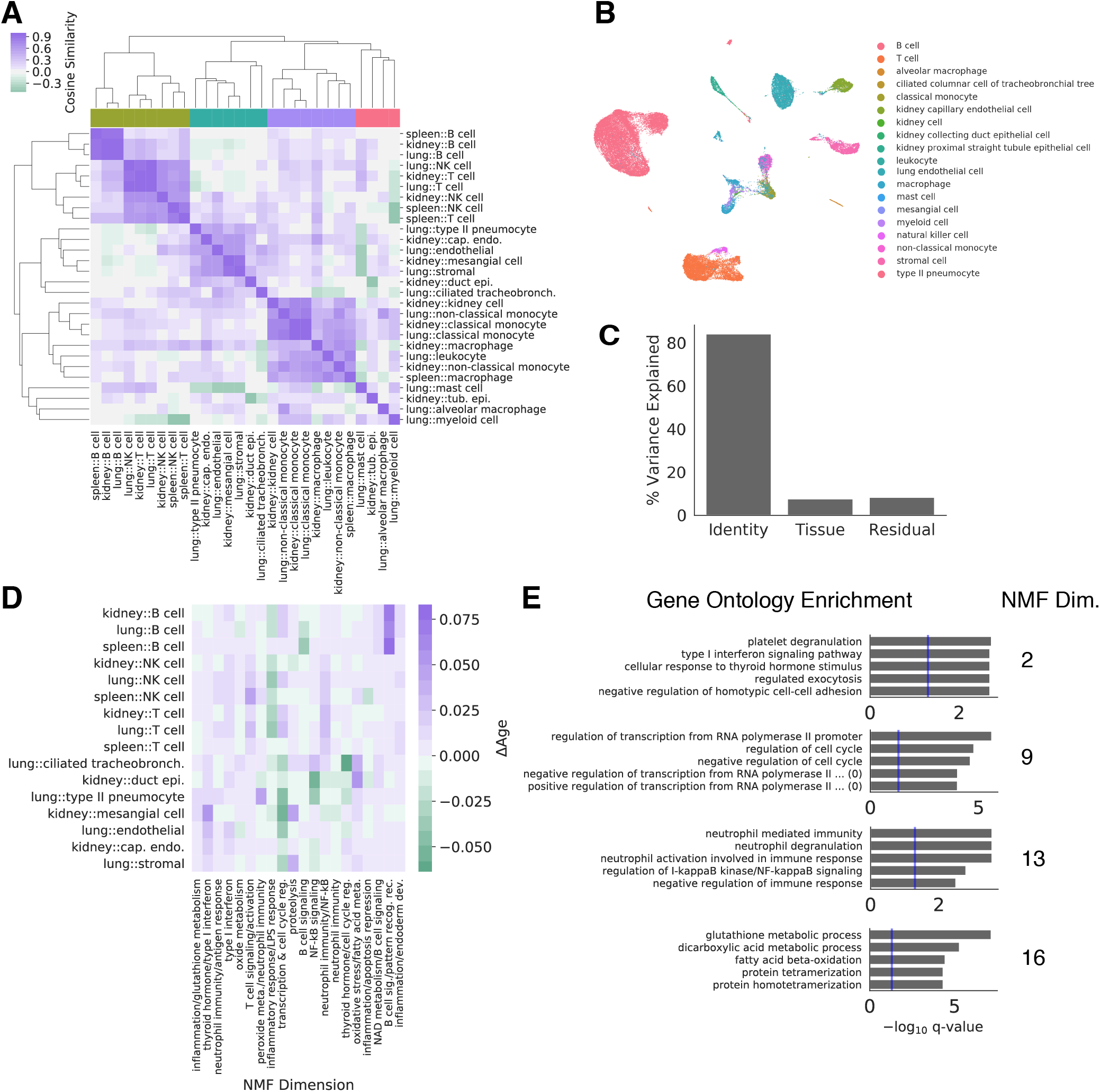
Cell identity and tissue environment influence aging trajectories. Aging trajectories were computed as the distance between young and old cell centroids in a non-negative low-rank embedding for each cell state. **(A)** Cosine similarities between the aging trajectories of each cell state in each tissue are compared in a heatmap. **(B)** A UMAP visualization derived from the 20-dimensional NMF embedding with cell types overlaid as colors. **(C)** Variation in the aging vectors of immune cell types found in all three tissues explained by cell type and tissue environment (ANOVA). **(D)** Heatmap visualization of the aging vectors for cell types. Endothelial cell types and lymphoid cell types identified by unbiased clustering presented in **(A)** are shown. Semantic descriptions of each embedding dimension derived from gene enrichment analysis are presented as column labels. Some expression programs show common changes with age across both groups of cell types, while others appear to be different between groups. **(E)** Gene enrichment analysis results for select dimensions of the NMF embedding.

We compare these trajectories using the cosine similarity. The cosine similarity is 1 if trajectories are in the same direction, 0 if they are orthogonal, and –1 if they are in opposite directions. Clustering by these cosine similarities, we find that qualitatively similar cell identities have similar aging trajectories, while dissimilar cell identities have orthogonal to dissimilar trajectories (Fig. 5A).

Examining the clustering partition, we find that cells segregate into roughly 4 clusters: endothelial and epithelial cells (blue), myeloid cells (purple), lymphocytes (green), and a remaining cluster with both myeloid and epithelial cell types (red). Cell identities from the kidney and lung are intermixed within the endothelial/epithelial cell cluster. Likewise, within the lymphocyte cluster B cells and T cells cluster more tightly by cell identity across tissues than by tissue of origin. This suggests that cell identity has a larger influence on aging trajectories than tissue environment.

To quantify the influence of both cell identity and tissue environment, we focused on the aging trajectories of immune cell types observed in all tissues (B cells, T cells, natural killer cells). For these cell types, we construct linear models and perform an analysis of variance (ANOVA) to determine the proportion of variation explained by cell identity and the proportion explained by tissue environment [77]. Consistent with qualitative observations, we find that cell identity explains a markedly larger fraction of variation than tissue environment (Fig. 5C). We also perform this analysis using a PCA embedding with the same result, indicating that this result is robust to our choice of embedding space (Supp. Fig. 13).

#### Endothelial cells and lymphocytes exhibit distinct aging trajectories

We next asked how endothelial/epithelial and lymphocyte aging trajectories differed. Using the clustering assignments from (Fig. 5A), we compared aging trajectories for each cell identity in the endothelial and lymphocyte clusters. We find that the two clusters show consistent differences in the magnitude of change within multiple NMF dimensions (Fig. 5D). Based on gene ontology enrichment within these dimensions, we find that endothelial/epithelial cell types show an increase in thyroid hormone and type I interferon signaling (Dimension 2, Fig. 5E) and oxidative/fatty acid metabolic processes (Dimension 16) relative to lymphocyte cell types. By contrast, lymphocytes show a larger increase in NF-*κ*B signaling and related immune response pathways (Dimension 13). We note that B cell activating associated pathways are increased with age in a B cell specific manner (Dimension 19). Both clusters show increases in NAD metabolism and B cell signaling pathways (Dimension 18), as well as T cell signaling and activation (Dimension 6). These results are consistent with literature observing changes in type I interferon activity [59] and thyroid hormone signaling with aging [36, 15, 94].

### Optimal transport analysis indicates that cell identity determines the magnitude of aging

Do some cell identities or tissue environments age more dramatically than others? To answer these questions, we estimated the magnitude of aging using optimal transport distances in an NMF embedding, as described above. Here, we utilize an embedding with 500 latent dimensions to capture more variation within the data (Supp. Fig. 11). Optimal transport is a technique for measuring distances between equally sized samples, such as probability distributions or discrete samples with the same number of elements. It was originally designed to calculate the minimal amount of earth that needed to be moved to convert a pile of earth into a fortification. Optimal transport distances have since been applied to a wide variety of tasks, including the analysis of single cell RNA-sequencing data [81].

The discrete optimal transport distance we apply here measures the minimum amount of change needed to make one group of cells match another. This metric capture differences in the covariance structure and modality of a cell population, in addition to differences in the population means (Supp. Fig. 14). We compare the distance between two cell populations by sampling *n* cells from each of them a number of times, and averaging the distance across the samples (see Methods). This bootstrap sampling scheme allows us to meet the equal sample size requirement for optimal transport distances, even when we observe different numbers of young and old cells.

Here, we compare the distance between young and old cells from each cell identity and tissue environment in the NMF embedding. For each cell identity, we make three distinct comparisons. We make a heterochronic comparison between young to old cells to estimate the magnitude of aging. This process is schematized in spleen B cells to provide intuition (Fig. 6A). We also make two isochronic comparisons, comparing young cells to young cells, and old cells to old cells (Fig. 6B). These isochronic comparisons serve as a null distribution, estimating the distance we would expect to see between random samples of cells in the absence of age-related change. In each comparison, we draw 300 samples, each containing *n* = 300 random cells (see Methods for sampling scheme used for low abundance cell identities). We normalize the heterochronic (Young-Old) comparison values for each cell identity by dividing by the mean of the larger isochronic distance.

**Figure 6:**
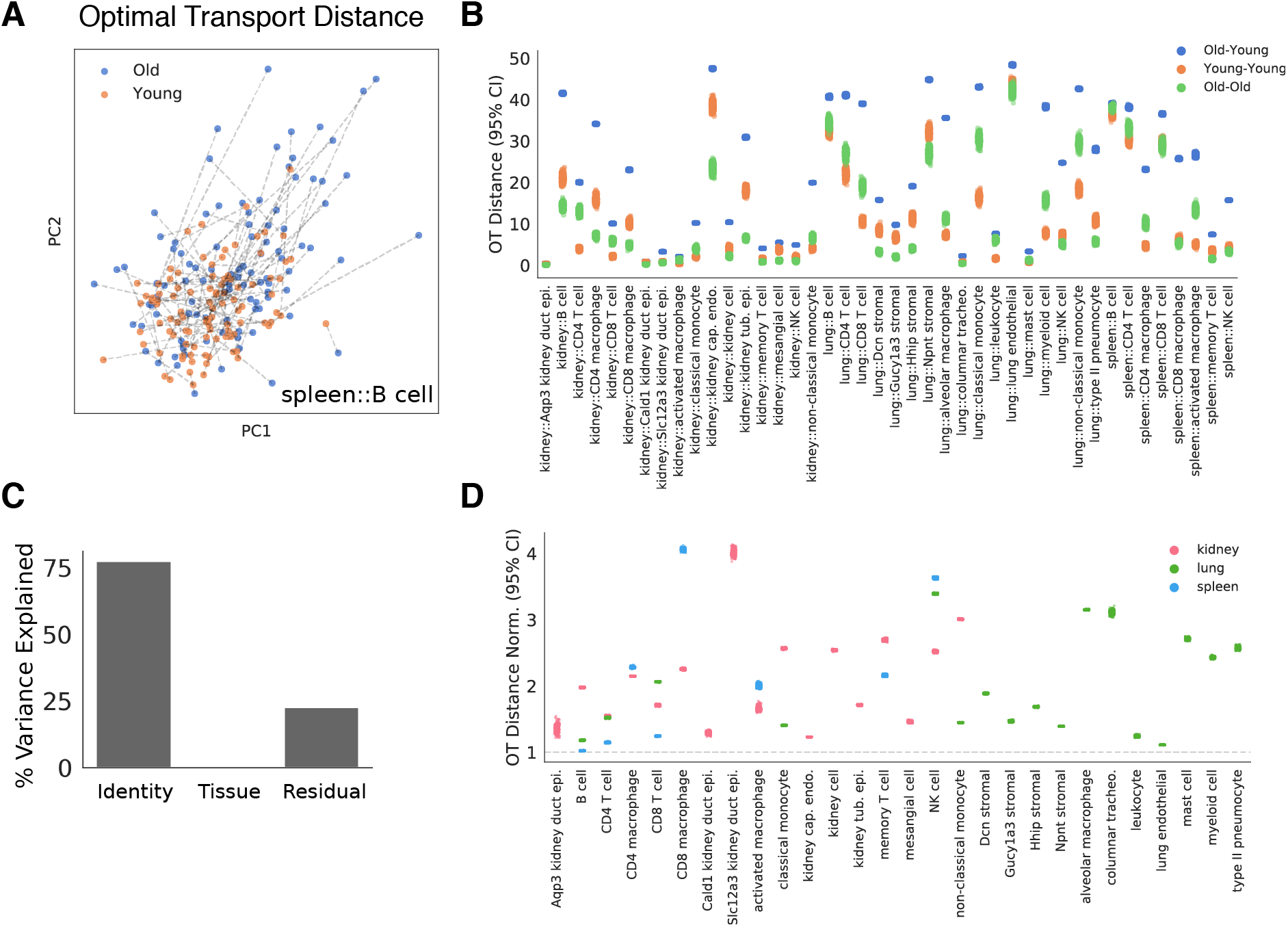
Optimal transport (OT) estimates the magnitude of aging across cell identities. To estimate a magnitude of age-related change in each cell population, we compute an OT distance between random samples of young and old cells in each cell identity. **(A)** Young and old spleen B cells are presented in a UMAP projection of the NMF embedding. Dashed lines overlaid indicate the globally optimal partners for each young and old cell, collectively representing the OT solution. The sum of distances along each dashed line is the OT distance. OT distances were computed independently for each cell identity across 300 random samples. **(B)** OT distances for each cell type in each tissue are presented. For each cell identity, we compute distances for heterochronic samples of cells (Old-Young) and isochronic samples of cells (Young-Young, Old-Old). The latter isochronic comparisons reflect negative controls. Distributions across *n* = 300 random samples each are presented for each of these comparisons. **(C)** Variance in normalized OT distances explained by cell identity and tissue environment (ANOVA). **(D)** Heterochronic OT distances for each cell identity. Each dot represents the normalized OT distance for a single sample. Values are normalized to the largest mean value of the isochronic negative controls. The grey dotted line marks a normalized distance of 1, which indicates that a heterochronic comparison shows similar distances to an isochronic comparison in that cell identity.

We interpret these normalized optimal transport distances as an estimate of the magnitude of age related change. We find a large diversity in aging magnitudes across the cell identities we observe. Some cell identities exhibit little more distance between young and old cells than between two isochronic groups of cells (normalized distance of ≈ 1), while others show 3 to 4 fold larger differences in heterochronic comparisons. Lung and kidney natural killer (NK) cells exhibit some of the highest optimal transport distances, while kidney and lung B cells and kidney capillary endothelial cells show some of the smallest distances.

To quantify the relative contributions of cell identity and tissue environment to aging magnitude, we use the same linear modeling and ANOVA approach as above. These models reveal that cell identity explains the majority of variation in aging magnitudes while tissue environment explains little variation (Fig. 6C). To confirm that these results are robust to the non-deterministic NMF embedding procedure, we compute these distances across 10 separate optimization runs of the NMF embedding and find that relative distances are preserved. We likewise compute Old-Young distances with a range of random sampling sizes and find distances are highly correlated across settings of this parameter (Supp. Fig. 15).

Perhaps surprisingly, we find that different cell states within the same cell type can have notable differences in aging magnitude. For instance, spleen CD8 macrophages exhibit a larger aging magnitude than spleen CD4 macrophages. Similarly, lung *Dcn* stromal cells exhibit a larger aging magnitude than other stromal cell counterparts. Across all three tissues, CD8 T cells exhibit a larger aging magnitude than CD4 T cells. These observations collectively indicate that cellular identity can have a notable impact on the magnitude of age-related change, such that even different cell states within the same cell type exhibit differences in aging magnitude. Tissue environments by contrast appear to have less influence on aging magnitude, suggesting that most difference in the magnitude of aging between tissues is driven by differences in cell identity composition.

## Discussion

Aging occurs across varied mammalian species, each composed of diverse cell types and states. While aging phenotypes are well catalogued at the organismal and tissue level, the cell identities that they comprise have been less explored. To understand the causes of aging, we wish to construct a causal network of molecular players and their relationships. If these players and relationships are dependent on cell identity, as evidence suggests for tissues [82, 43, 12, 72], an accurate representation of the causal network requires resolution of the organismal building blocks – individual cells.

Here, we use single cell RNA-seq to investigate aging across a diverse set of murine cell identities in three tissues. Tissues were collected in synchrony from each animal, allowing for direct comparisons between cell identities within and across tissues. Few synchronized cross-tissue data sets investigating aging phenotypes are available, and we believe these data will be informative to the aging research community.

We find that cell identities exhibit unique differentially expressed genes with aging, consistent with previous reports of cell identity specific aging phenotypes [4]. Similar cell types (e.g. kidney capillary endothelial cells & lung endothelial cells) exhibit broadly similar aging trajectories across tissues, while distinct cell types from the same tissue (e.g. B cells & type II pneumocytes in the lung) have dissimilar trajectories. This suggests that cell identity and aging trajectory are coupled, and distinct cell types may be undergoing independent aging processes. Consistent with this notion, cell identity explains the majority of variation in these aging trajectories. Tissue environment explains a lesser amount variation, but its influence is also present in our data in small sets of unique differentially expressed genes in the same cell identity across tissues. Collectively, our results indicate that the molecular manifestations of aging differ between cell identities and tissue environments. A causal network for aging phenotypes must therefore be conditioned on both cell identity and environment to accurately reflect biology.

Although most changes are unique to tissues or cell types, we did identify a shared core of differentially expressed genes with age. This core is characterized by decreased expression of genes involved in SRP-dependent protein translation and protein targeting to the ER and increased expression of genes involved in inflammation. Decreased protein translation and targeting to the ER has previously been associated with replicative age in *S. cerevisiea* [40] and is consistent with associations between global protein translation and aging [87]. Likewise, increased inflammatory pathway expression has been broadly observed in studies of aging tissues [12, 70, 78, 3]. Observation of these changes across many cell identities and tissue environments suggests they are consistent molecular players in the causal network of aging, less dependent on context than others. We caution that consistency and causality are not intertwined, and functional studies modulating these pathways in multiple cell identities are necessary to establish causal links to other aging phenotypes.

We present an optimal transport metric to estimate the magnitude of age-related change between two cell populations. Using this approach, cell identities exhibit multi-fold differences in aging magnitude, and cell identity again explains the majority of variation in aging magnitudes. Our results are conceptually consistent with previous reports of dramatic transcriptional changes with age in some cell identities and more subtle changes in others [46, 47, 53, 28]. Directly comparing the magnitude of aging in this manner suggests that some cell identities may exhibit more dramatic functional decline with aging than others. However, the magnitude of transcriptional change with aging may not necessarily reflect the degree of functional decline in all cases. Future comparison between identities with high and low aging magnitudes may reveal some cellular “strategies” that moderate age-related change.

Tissue level profiling of aging tissues has suggested that cell type composition may change with age [12, 70, 78]. Increased immune gene expression in multiple old tissues has specifically suggested increased infiltration of immune cells. Our data are consistent with this notion, as we observe an increased frequency of immune cell types in both the kidney and lung. However, these changes are difficult to confirm using only cell type counts produced after single cell isolation. Age-related changes in a tissue (i.e. altered extracellular matrix composition [4]) may lead to preferential isolation of one cell type relative to another, even if the underlying cell type proportions do not change. To confirm this phenotype, immunohistology studies quantifying the number of immune cells in these aging tissues are necessary. We also observed that inflammatory pathways are commonly upregulated, even in non-immune cell identities, suggesting that changes in immune cell proportions alone do not entirely explain the increased inflammatory pathway activity in aging tissues.

Previous studies reported that multiple metrics of transcriptional variability increase with aging [67, 34]. Two measures have predominated in the literature – (1) gene expression variance measures which focus on the mean “over-dispersion” across genes and (2) cell-cell heterogeneity measures which focus on the mean distance of individual cells to the center of the cell population. While ostensibly similar, each measure provides insight into a different aspect of biology.

Gene expression variance does not weight genes by expression level, such that changes are most likely when many genes increase or decrease variance. Biologically, changes in this measure may therefore reflect global changes in transcriptional noise [33, 76], which has been reported to influence cell fate decisions [96, 24, 41, 9]. By contrast, cell-cell heterogeneity measures consider the absolute distance of each cell from the population center, such that changes in heterogeneity can be driven by a handful of moderate- or high-expression genes. These measures may better reflect the phenotypic variance across cells in a population, as associated with the classical bet-hedging phenomenon [26, 85, 10]. We find that changes in each of these measures depend on cell identity, such that neither increased or decreased variability was observed universally across identities. These results are consistent with previous reports of aging altering gene expression variance and cell-cell heterogeneity in a cell identity dependent manner [4].

## Conclusions

Single cell RNA-seq of young and old cells from three separate mouse tissues has revealed common and cell identity specific aspects of aging. We find that multiple metrics of transcriptional variation change with age, and that these changes are cell identity dependent. Cell identities exhibit unique gene expression changes with age, but we also identify a common set of protein translation and inflammatory pathway changes across many cell identities. Leveraging matrix factorization techniques, we define trajectories of aging and attribute the majority of variation in these trajectories to cell identity. Using optimal transport, we compute a magnitude of aging and find that observe multi-fold differences in this magnitude between cell identities. We again find that cell identity rather than tissue environment explains the majority this variation in magnitude. Collectively, these results highlight the influence of cell identity on the aging process and the importance of measuring aging phenotypes with cellular resolution.

## Methods

### Animals

Young (29-34 weeks old) and old (88-93 weeks old) male C57Bl/6 mice were used for all experiments. Mice were housed communally with a standard 12 hour dark cycle and fed *ad libitum*. Euthanization was performed by administration of carbon dioxide at a controlled flow rate. Animals were euthanized at the same time each day to minimize circadian variation, shortly after the beginning of the light cycle. Kidneys, lungs, and spleen were collected from each experimental animal and weighed. Downstream cell isolation for each tissue proceeded immediately.

### Cell Isolation

We removed each tissue and washed each in HBSS then dissected the tissue into small pieces using a razor blade.

#### Kidney and Lung

We incubated dissected kidney tissue in 1.48 U/mL Liberase DL enzyme mixture (Roche) in a total volume of 7 mL DMEM at 37°C for 20 minutes in a 50 mL conical tube shaking at 200 rpm. We incubated dissected lung tissue in 1.48 U/mL Liberase TM (Roche) and 200 U/mL DNase I (Roche) in a total volume of 7 mL DMEM at 37°C for 30 minutes in a 50 mL conical tube shaking at 200 rpm. Tissue was further mixed using a 10 mL pipette tip and 40 mL of DMEM (2% FBS) was added to stop digestion. We sequentially pipetted cells through 100 *μ*m, 70 *μ*m, and 40 *μ*m filters then peletted cell suspensions by centrifugation at 500 × *g* for 10 minutes.

After centrifugation, we treated cells with ACK Lysing Buffer (ThermoFischer) for 5 minutes at room temperature. We centrifuged ACK treated cells, discared the supernatant, and repeated ACK treatment. We subsequently washed cells in DMEM (2% FBS) and peletted by centrifugation. We removed debris from samples using the Miltenyi Debris Removal Solution. We added 6.2 mL cold PBS and 1.8 mL Debris Removal Solution to cell pelettes and resuspended by pipetting. We added 4 mL cold PBS to the top of the mixed solution and centrifuged samples at 4°C, 3000 × *g* for 10 minutes. We washed samples in 10 mL cold PBS and centrifuged at 4°C, 1000 × *g* for 10 minutes, then resuspended in PBS with 2% FBS. We counted cells using a TC20 Cell Counter (Bio-Rad) and diluted cells to a concentration of 10^6^ cells/mL. We then proceeded to single cell library preparation.

#### Spleen

We used a syringe plunger to further fragment the spleen tissue after razor blade dissection. We mechanically dissociated tissue fragments by pipetting in RPMI cell culture media (2% FBS). We subsequently forced tissue fragments through 100 and 40*μ*m filters. After centrifugation at 500 × *g* for 10 minutes, we treated cells with ACK Lysing Buffer (ThermoFischer) for 5 minutes at room temperature. We washed cells in RPMI (2% FBS) and pelleted by centrifugation. We counted cells using a TC20 Cell Counter (Bio-Rad) and diluted cells to a concentration of 10^6^ cells/mL then proceeded to single cell library preparation.

### Single Cell RNA Sequencing Experiments

We prepared libraries (individual lanes on the 10X Chromium) with the 10X Single Cell 3’ v2 kit using 6,000 cells per lane on the 10X Chromium microfluidics device. We sequenced libraries at a target depth of 50 million reads/sample on an Illumina HiSeq 4000. For each sample, we performed two technical replicates by preparing two separate libraries from the same cell suspension across two channels of the Chromium microfluidic device.

### Read Alignment and Gene Expression Quantification

We aligned reads to the `mm10` reference genome obtained from ENSEMBL. We used gene annotations from the GENCODE vM20 release with slight modification. We have replaced annotations for lincRNAs *Gm42418* and *AY036118* with a single contiguous gene annotation for the rRNA element *Rn45s*. This locus harbors an *Rn45s* repeat as reflected in RefSeq, such that contaminating 18S rRNA in our library preparations may lead to inflated expression counts for these lincRNAs.

We performed alignment to this amended reference using 10X `cellranger 3.0.2`, which employs the `STAR` sequence aligner [30]. We determined gene expression counts using unique molecular identifiers (UMIs) for each cell barcode-gene combination. Following alignment, we filtered cell barcodes to identify those which contain cells using the approach implemented in `cellranger 3.0.2`, and only these barcodes were considered for downstream analysis. The output of this analysis is a Cells × Genes matrix, where each element *i, j* represents the number of UMIs mapping to gene *j* in cell *i*.

### Quality Control

We removed libraries which contained a low sequencing depth (< 10 million reads) or very few cells detected (< 500 cells) from subsequent analyses as likely experimental errors. Based on this metric, only one technical replicate failed QC, so all tissues from all animals in the experimental design are represented in our downstream analysis. We leveraged the `scanpy` toolkit [99] in subsequent analyses. We quantified the total number of UMIs, total number of genes detected, fraction of UMIs mapping to the mitochondrial genome, and fraction of reads mapping to the *Rn45s* repeat annotation for each cell. We filtered cells with fewer than 1000 UMIs or fewer than 500 unique genes detected, as described for the *Tabula Muris* [80]. Additionally, we filtered out cells with a high fraction of reads mapping to the mitochondrial genome (> 10%) or the *Rn45s* repeat (> 0.5%) as likely dead cells.

### Dimensionality Reduction and Embedding

We normalized the Cells × Genes matrix by the total number of UMIs in each cell and scaled by 10^6^ to yield Counts Per Million (CPM). We natural log transformed this matrix after the addition of a pseudocount of 1 to avoid undefined values.

To enable unsupervised clustering and cell type identification, we perform dimensionality reduction with principal component analysis (PCA) to the combined set of samples for each tissue. First, we identify a set of highly variable genes within the tissue based on overdispersion as described previously [79]. Briefly, we computed a “normalized dispersion” score for each gene by binning genes with similar mean expression levels, and subtracting the mean dispersion (variance/mean) within a bin from the dispersion score for each gene in the bin. The residual values represent the dispersion after accounting for the mean:variance relationship using this binning scheme. We set the following minimum criteria for the selection of genes as highly variable: minimum overdispersion of 0.5, minimum mean expression of 0.5 ln(CPM + 1), maximum mean expression of 7 ln(CPM + 1). After subsetting to this set of highly variable genes, we centered expression values to a mean of 0 and unit variance. We used these scaled genes as input features for PCA. Once embedded in this PCA space, we construct a nearest neighbor graph identifying the *k* = 15 nearest neighbors for each cell. We derived UMAP embeddings presented for visualization from this nearest neighbor graph using a minimum distance of 0.5 and a spread of 1.0 [11].

### Clustering and Cell Type Identification

Louvain community detection [79, 19] was applied to the nearest neighbor graph constructed in PCA space to define a cluster partition. To infer cell types, we trained a neural network classifier to predict cell ontology classes given single cell RNA-seq expression information.

This classifier is a fully connected neural network with four hidden layers each paired to a rectified linear unit activation, dropout layer, and batch normalization. Each hidden layer has 1024 units and the dropout probability on each layer is set to *p* = 0.3. We apply a softmax activation to the final layer and utilize cross entropy as an objective function for training. During training, we perform class balancing with a mixture of over- and under-sampling. Classes (cell types) with fewer than 128 examples are oversampled, while classes with more examples are undersampled. Optimization is performed using the Adagrad optimizer [31].

As a training set, we utilized the recently published *Tabula Muris* compendium which provides expert cell type annotations in the mouse [80]. In addition to these annotations, we manually added cell state annotations to the Tabula Muris data to provide a level of granularity below cell ontology classes (Supp. Fig. 1). Example cell states include categorizing T cells into CD4 T cell and CD8 T cell subgroups, as well as the addition of subgroup labels to heterogeneous cell types such as lung stromal cells in the *Tabula Muris*. We name these cell states which do not have canonical names based on the expression of a prominent marker gene. For instance, we refer to a *Gucy1a3*+ subset of lung stromal cells as *Gucy1a3* lung stromal cells.

We first trained a classifier for each tissue individually, yielding a set of tissue-specific classifiers that have no knowledge of cell types outside the training tissue. We also trained a tissue independent classifier by training on all cell types present in the Tabula Muris simultaneously.

We first inferred cell types for the lung and spleen using a tissue-specific classification model trained to predict cell ontology classes. We classified kidney cells with a tissue independent model to provide further resolution of immune cell types not annotated in kidney cells of the training set. We subsequently inferred subtypes using a model trained to predict our added subtype annotations. We chose subtypes as the most likely subtype allowed within our defined hierarchy. For instance, we chose the subtype of a T cell as the most likely subtype among the “CD4”, “CD8”, and “memory” subtypes. We used subtype classification models trained on the spleen to predict T cell and macrophage subtypes in all tissues, as the training set contained insufficient cell numbers to perform subtype annotation in other tissues. After direct inference of types and subtypes, we refined cell type information by using a *k*-nearest neighbors smoothing approach. Here, we chose *k* ∈ [30, 100] empirically depending on the tissue context.

### Differential Cell Type Proportion Analysis

To determine if cell type proportions differed between old and young animals, we performed an additive log ratio (ALR) transform on the observed cell type frequencies and assessed significant changes for each cell type using a *t*-test. Within a given cell type, we performed a *χ*^2^-test of the Age × Cell State contingency table to determine if the proportions of cell states change with age.

### Differential Variability Analysis

We measured differences in transcriptional variation between young and old animals in two distinct ways.

1. The first method evaluates changes in the variability of each gene between young and old animals, and attempts to identify a shift in the distribution of gene-wise variation. We assessed gene-specific variability by measuring the “overdispersion” of each gene. We defined overdispersion as the residual between a gene’s observed dispersion and the expected dispersion based on the gene’s mean expression value. We computed overdispersion values using the “difference from the median” (DM) method, as introduced previously [53]. We restricted DM calculation to genes with a mean expression value greater than 35 CPM to reduce the influence of poorly measured genes. To determine if the distribution of overdispersion values is significantly changed across ages, we employ the Wilcoxon Rank Sums test for a difference in means. We controlled the False Discovery Rate (FDR) to *α* = 0.05 with the Benjamini-Hochberg procedure. We performed DM analysis for each cell state in each tissue in our data set separately.
2. The second method we employ evaluates cell-cell heterogeneity based on the Euclidean distance between cells in expression space, as introduced previously [34]. For each cell state in each tissue, we compute the centroid of the cell state in gene expression space. We computed the distance from each cell to this centroid as a metric of cell-cell heterogeneity within each cell state. We employed the Wilcoxon Rank Sums test with the Benjamini-Hochberg procedure [13] as before to determine if this cell-cell heterogeneity metric is significantly different across ages.

### Differential Expression Analysis

We computed differentially expressed genes between two groups of cells A and B using the Wilcoxon Rank Sums test with the Benjamini-Hochberg procedure for FDR control. We performed differential expression between young and old cells within each cell state in each tissue independently. In addition to comparing mean expression values with the Rank Sums test, we also computed the proportion of cells expressing each gene in each group. Cells with >= 1 UMIs for a given gene are considered to be expressing the gene, whereas cells with 0 UMIs are considered to not be expressing the gene.

#### Identification of Common and Unique Differentially Expressed Genes

To identify common differentially expressed genes with age across cell identities, we computed for each gene the number of cell identities in which that gene is significantly differentially expressed in the same direction. Note that we counted each cell identity as a single entry in this score, regardless of how many tissues it appears in. We then selected genes that are differentially expressed in the same direction in *k* >= 5 cell identities and consider this gene set to be “commonly differentially expressed” with age.

To identify genes which are uniquely differentially expressed between two cell identities **A** and *B*, we computed genes which are significantly differentially expressed in *A* and are (1) not-significantly differentially expressed in *B* and (2) have a log_2_ fold-change < 0.1 in *B*.

### Gene Ontology Enrichment Analysis

We used `Enrichr` [54] to perform gene set enrichment analysis against the Gene Ontology Biological Process (2018) gene set collection. We also used the MSigDB Hallmark Gene Sets [61], for which we computed enrichment scores using Fisher’s exact test. In both cases, we corrected for multiple hypothesis testing using the Benjamini-Hochberg procedure.

### Cell Cycle Scoring

Cell cycle activity was estimated by scoring the expression of a set of S phase associated and G_2_/M phase associated genes, as shown previously [90] and as implement in `Seurat` [79]. Briefly, a set of genes associated with each of these phases was derived from single cell RNA-sequencing in 293T and 3T3 cell lines. For each cell, the sum of the expression of these phase-associated genes is computed. As a null distribution, a set of genes are selected from the set difference of the observed genes and the phase-associated genes. These null genes are selected by binning the sample genes into 24 equally sized bins by mean expression, then randomly sampling 100 null genes per bin per phase-associated gene with replacement. The difference between the mean of the phase-associated genes and the selected null genes is considered the module score.

### Analysis of Variance in Transcriptional Space

To determine the proportion of variance in transcriptional space (gene-wise UMI counts, NMF embedding dimensions) explained by experimental factors in our data, we use the linear modeling approach of Robinson *et. al*. [77]. Briefly, we fit a linear model for each dimension of the relevant transcriptional space (i.e. gene, NMF dimension) of the form

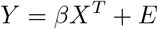

where *Y* is a Feature × Cell matrix of observed transcriptional features, *X* is a Samples × Parameters design matrix containing *p* experimental parameters, *β* is a Feature × Parameters coefficients matrix, and *E* is a Feature × Cell matrix of residuals. We calculated the proportion of variance explained by each experimental parameter for each gene by ANOVA. To determine the total proportion of variance explained by a factor, we simply sum the sum of squares for each parameter *p* across features for each parameter and divide by the summed total sum of squares:

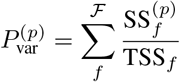

where *f* is a feature in the set of features 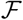.

### Non-negative Matrix Factorization Embedding

We perform non-negative matrix factorization using a standard multiplicative update optimization and random initialization [57], as implemented in the `nimfa` package [103]. We performed NMF optimization using all cells observed across all three tissues after ln(CPM + 1) normalization. The NMF embedding was fit to a set of highly variable genes, identified as described above. We chose a rank *k* = 20 for aging vector by selecting the “knee” in a plot of Rank vs. Explained Variance. We also utilize an NMF embedding of rank *k* = 500 for optimal transport distance calculation to capture more variation in the data, also chosen based on a later “knee‘ in the Rank vs. Explained Variance curve.

#### NMF Embedding Interpretation

To assign semantic meaning to each dimension of the embedding, we first identify genes associated with each dimension by performing Otsu thresholding [73] on log-transformed gene loadings. From the set of associated genes (loadings above the threshold), we perform gene ontology enrichment analysis with the Biological Process Gene Ontology database. We empirically name a consensus “program” for each dimension based on the enriched gene sets.

### Aging Trajectory Calculation

We compute aging trajectories for each cell identity/tissue environment combination individually. For each cell identity, we compute the centroid *c* of the young cells and old cells in the NMF embedding and compute the vector from the young cells to the old cells 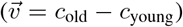.

### Optimal Transport Estimation of Aging Magnitude

We use a discrete optimal transport (OT) distance to estimate the magnitude of difference between two cell populations *A* and *B*. Due to the conservation of mass assumption in the optimal transport formulation, we take random samples of the same size *n* from each population for comparison. We compute the OT distance as minimum cost solution to the linear sum assignment problem, solved using the Munkres algorithm [68]. For each comparison of populations *A* and *B*, we perform 300 random samples of size *n* = 300 cells and compute the average to reduce the variability inherent in this stochastic sampling approach. In the data set examined here, multiple populations have < 300 cells for sampling. In this circumstance, we set *n* = 0.8min(∥*A*∥, ∥*B*∥) and use the same repeated sampling approach as above.

To estimate the “magnitude of aging,” we compute OT distances for three distinct comparisons. We make a heterochronic comparison of young cells to old cells as a measure of the difference between these populations (Old-Young comparison). As negative controls, we compute both isochronic comparisons, measuring distances between random samples from pool of young cells (Young-Young comparison), and measuring distances between samples from the pool of old cells (Old-Old comparison). The isochronic comparisons serve as an estimate of the distance we would expect between random samples simply due to heterogeneity within the population.

We normalize the heterochronic Old-Young comparison by dividing these measurements by the mean of the largest isochronic distance (i.e. if Young-Young distance is larger than Old-Old, we divide by the Young-Young mean, and vice-versa). This normalization scheme is conservative, using the upper bound estimate of differences caused by cell-cell heterogeneity as our baseline for noting an effect due to aging. Following normalization, we therefore interpret Old-Young OT distances > 1 as indicative of differences caused by aging and interpret a larger value of this normalized distance as reflecting a larger magnitude of age-related change. To ensure these distance estimates are robust to different optimizations of the NMF embedding, we perform NMF optimization using 10 distinct random initializations and compute OT distances in each of these embedding spaces.

### Software Tools

We leveraged GNU parallel [66], the scipy computing environment [71], and the Seaborn plotting package for several analyses [95].

### Data Availability

Experimental data have been submitted to the Gene Expression Omnibus [32]. Additionally, we provide access to relevant data and analysis code on our website: https://mca.research.calicolabs.com/.

## Competing interests

The authors are current and former employees of Calico Life Sciences.

## Author’s contributions

JCK performed analysis and wrote the paper. LP conceived the study and performed experiments. NDR performed analysis. DGH performed analysis and provided experimental support. DRK supervised research and wrote the paper. AZR conceived the study, supervised research, and performed experiments.

## Acknowledgements

We thank the Calico Physiology group, including Chunlian Zhang, Adam Kennedy, and Ganesh Kolumam for animal experiment support. We thank the Calico Genomics group led by Margaret Roy for assistance with library preparation and sequencing. We thank Allon Klein and Daniel Gottschling for helpful discussions and support.

## Supplementary Information

**Figure S1:**
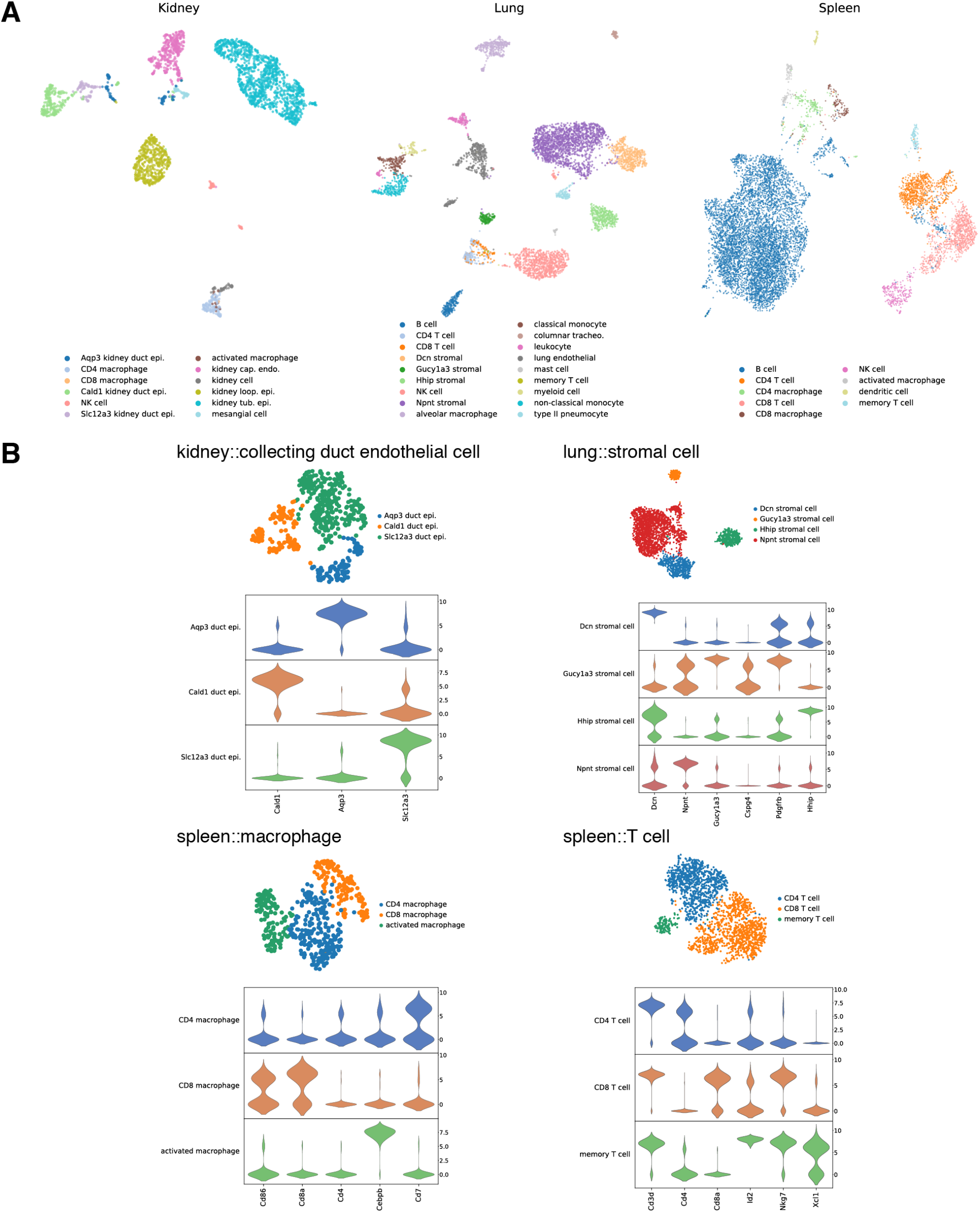
Manual annotation of cell states in the Tabula Muris. **(A)** UMAP projections of the Tabula Muris data with our manual cell state annotations overlaid as colors. We annotate cell states where there are known canonical states within a cell type (i.e. CD4 and CD8 T cells) and where we observe substructure within a cell type in the *Tabula Muris* data (as in the lung stromal cells). **(B)** UMAP projections of each cell type where we provide manual cell state annotations. The expression of marker genes that guided our cell state annotations is shown below each UMAP projection.

**Figure S2:**
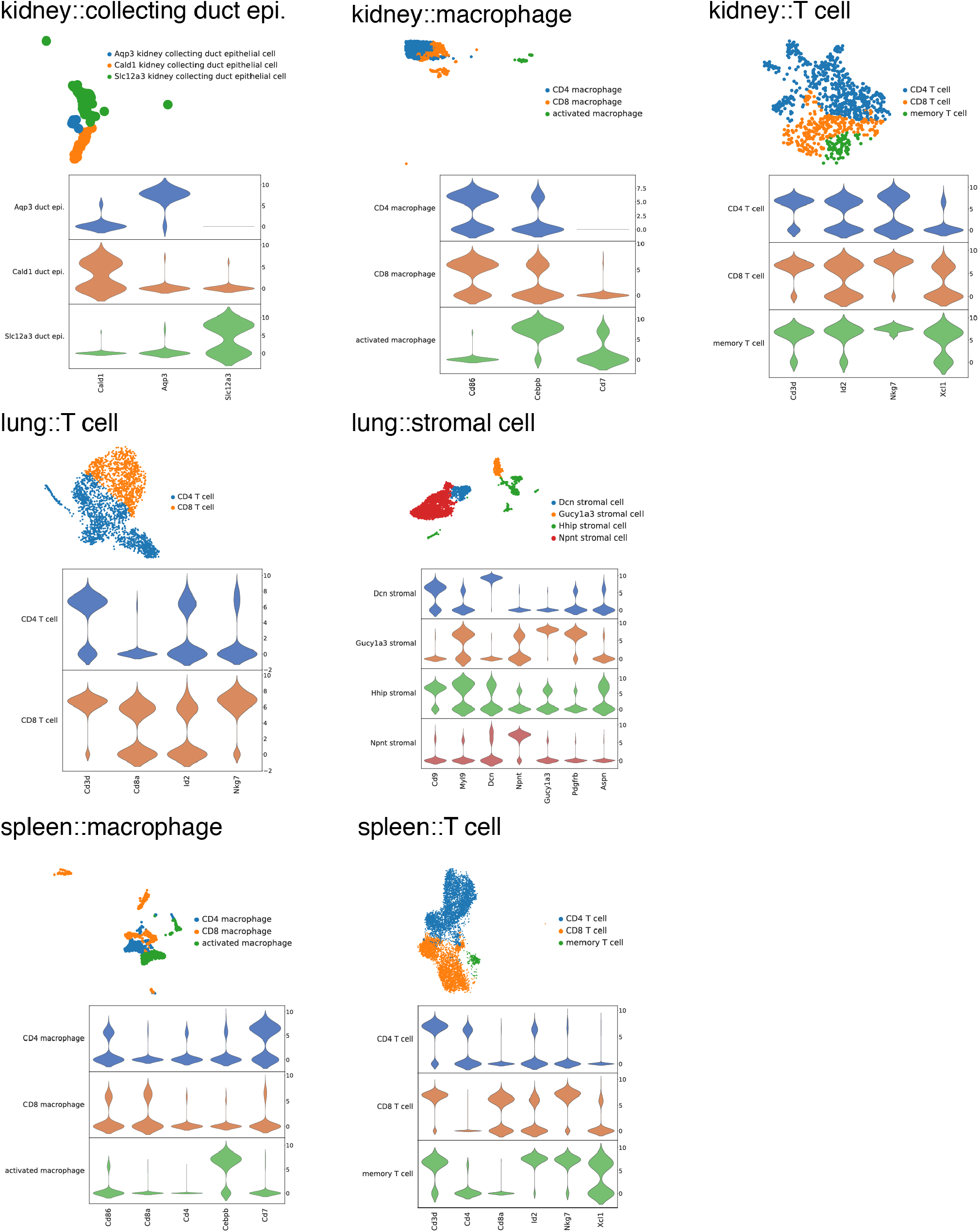
Cell state annotations in our data derived from neural networks. We trained deep neural networks to classify cell states within individual cell types using our manual annotations of the *Tabula Muris* (Supp. Fig. 1). UMAP projections of each cell type are presented with cell state annotations overlaid as color labels. Cell states are enriched for corresponding marker genes, as in the *Tabula Muris*, presented as violins below each UMAP projection.

**Figure S3:**
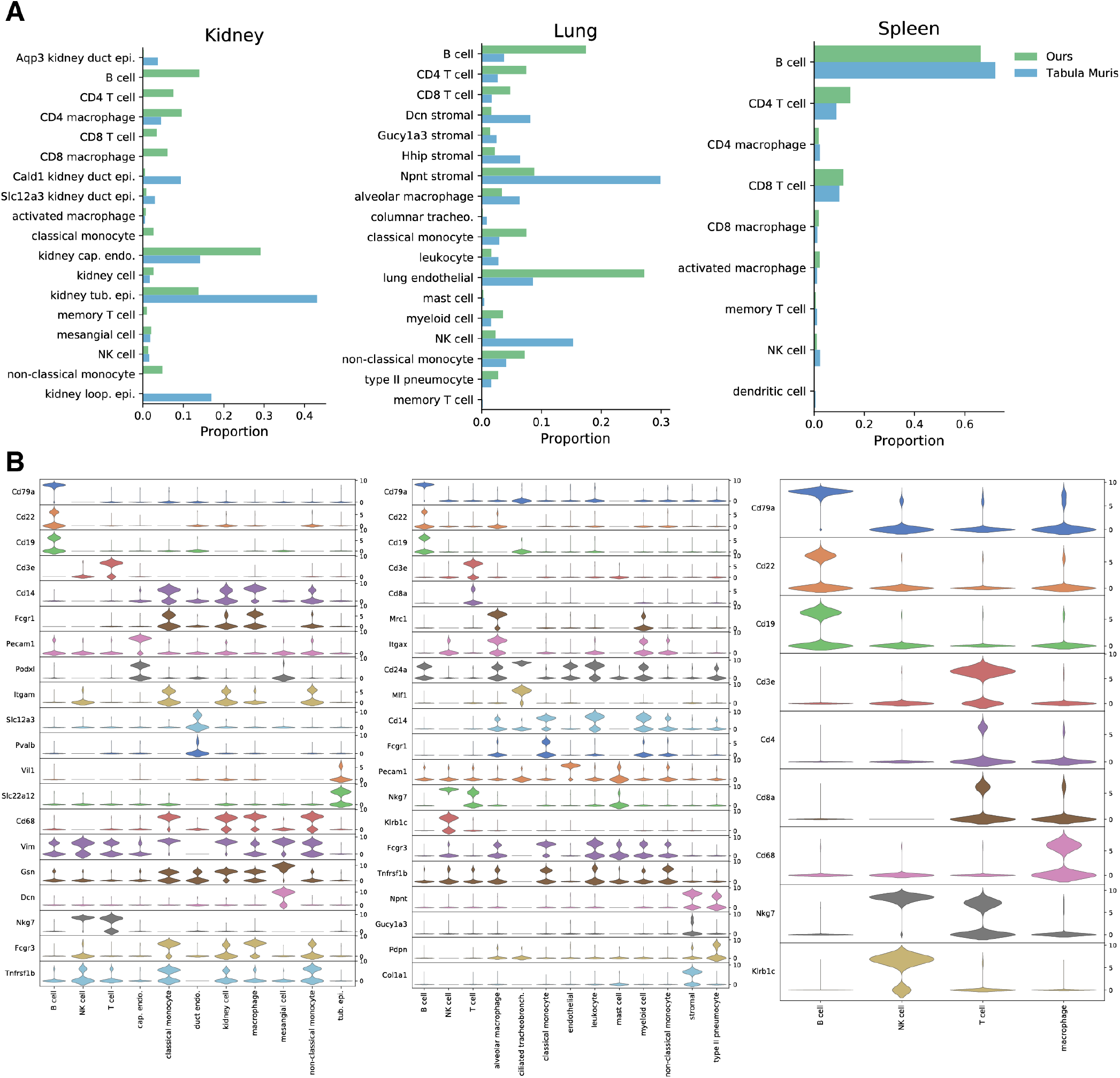
Comparison of our cell state proportions and markers to the Tabula Muris. **(A)** Cell state proportions from our dataset (green) compared to proportions observed in the Tabula Muris (blue). Notable differences include a higher proportion of immune cells in our data set and the absence of kidney loop of Henle epithelial cells. These differences may be the result of intentional experimental differences (perfusion vs. no perfusion), animal ages (Tabula Muris animals are younger than our young animals), and laboratory-to-laboratory differences in isolation technique. **(B)** Expression of marker genes across cell states we identify in our data. Marker genes are either taken directly from those suggested by the Tabula Muris, or from marker genes we identify in their data using our differential expression procedure.

**Figure S4:**
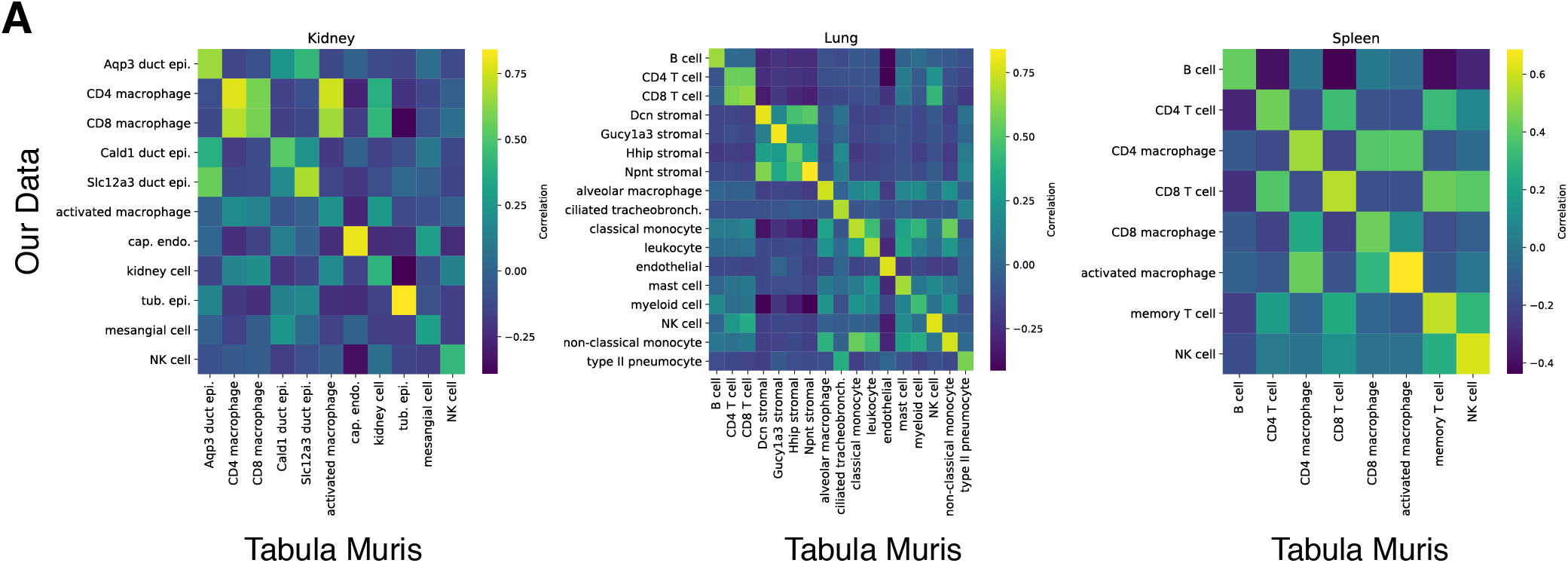
Correlation of cell identities between our data and the *Tabula Muris*. **(A)** Mean expression vectors for each cell identity were computed in our data and the *Tabula Muris* as the mean expression of each gene across cells in a given identity. We computed correlations between these mean expression vectors for all identities between the two data sets. Visualizing these correlations as a heatmap, we find that identities in our data are most similar to corresponding identities in the *Tabula Muris*.

**Figure S5:**
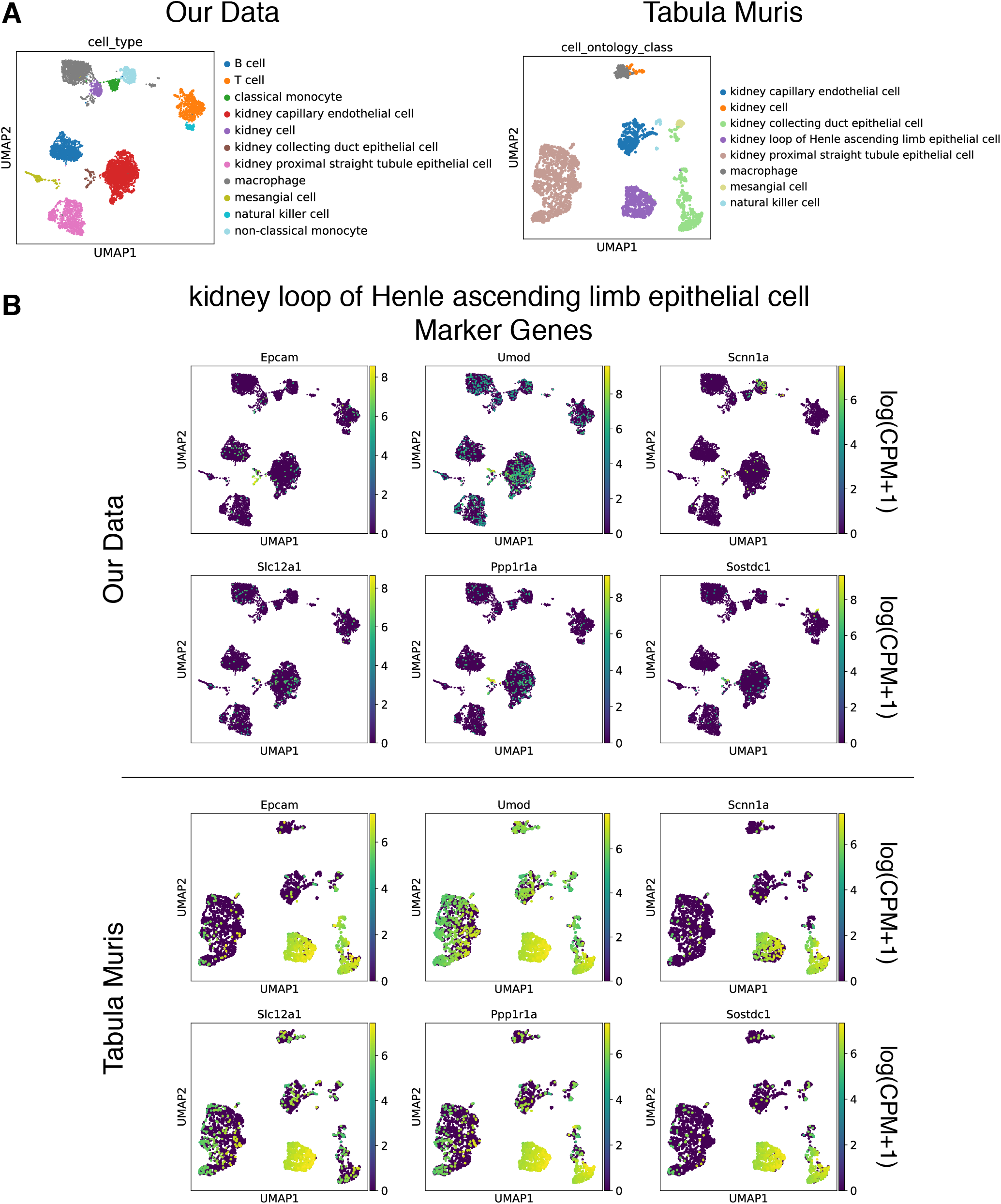
Visualization of kidney loop of Henle epithelial cell markers from the Tabula Muris indicates these cells are not present in our data. **(A)** UMAP projections of our kidney data and kidney 10X kidney data from the Tabula Muris. Note the absence of kidney loop of Henle epithelial cells in our data set. **(B)** Expression of marker genes for kidney loop of Henle epithelial cells in our dataset and the Tabula Muris. We find no coherent group of kidney loop of Henle cells in our data based on these markers.

**Figure S6:**
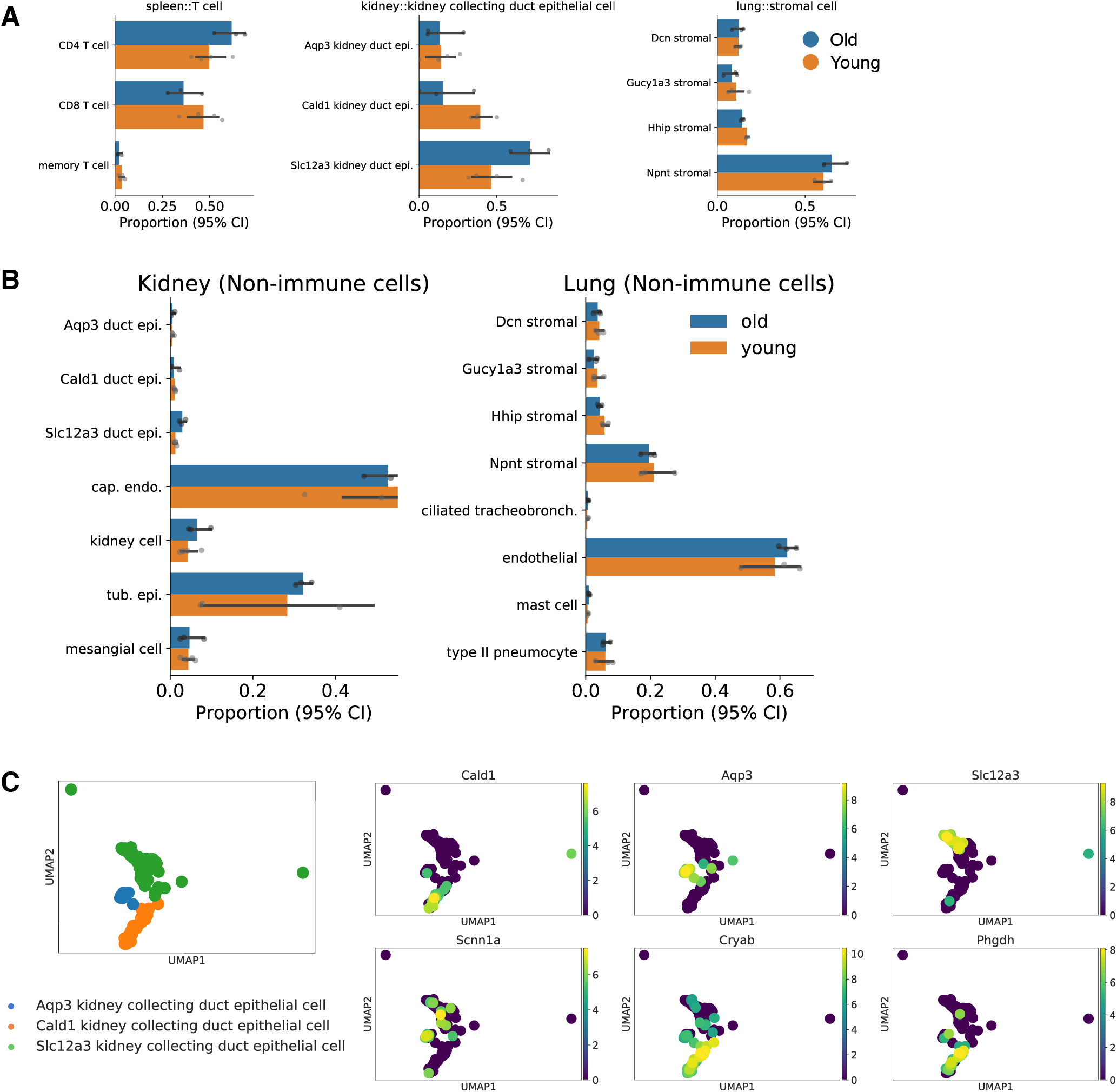
Cell state proportions shift with age. **(A)** Cell subtype proportions with age for individual cell types. Kidney collecting duct epithelial cells (left) show a significant shift in the subtype distribution with age (Chi-squared contingency table, *p* < 0.05). By contrast, lung stromal cells with a similarly complex cell type composition do not change with age (center). In the spleen, we detect a well known shift in the T cell subtypes with age. Old animals exhibit a lower proportion of CD8 T cells (Chi-squared contingency table, *p* < 0.05). **(B)** Cell subtype proportions with age for all non-immune cell types in the kidney and lung. When immune cells are removed, we find that most cell types in both tissues have similar proportions between young and old animals. **(C)** *Aqp3* and *Slc12a3* kidney collecting duct epithelilial cells co-express *Scnn1a* (epi. sodium channel ENaC) suggesting they are principal cells of the collecting duct. *Cald1* duct epithelial cells also express *Phgdh* and *Cryab*, consistent with reports of a subpopulation in the literature with unknown function.

**Figure S7:**
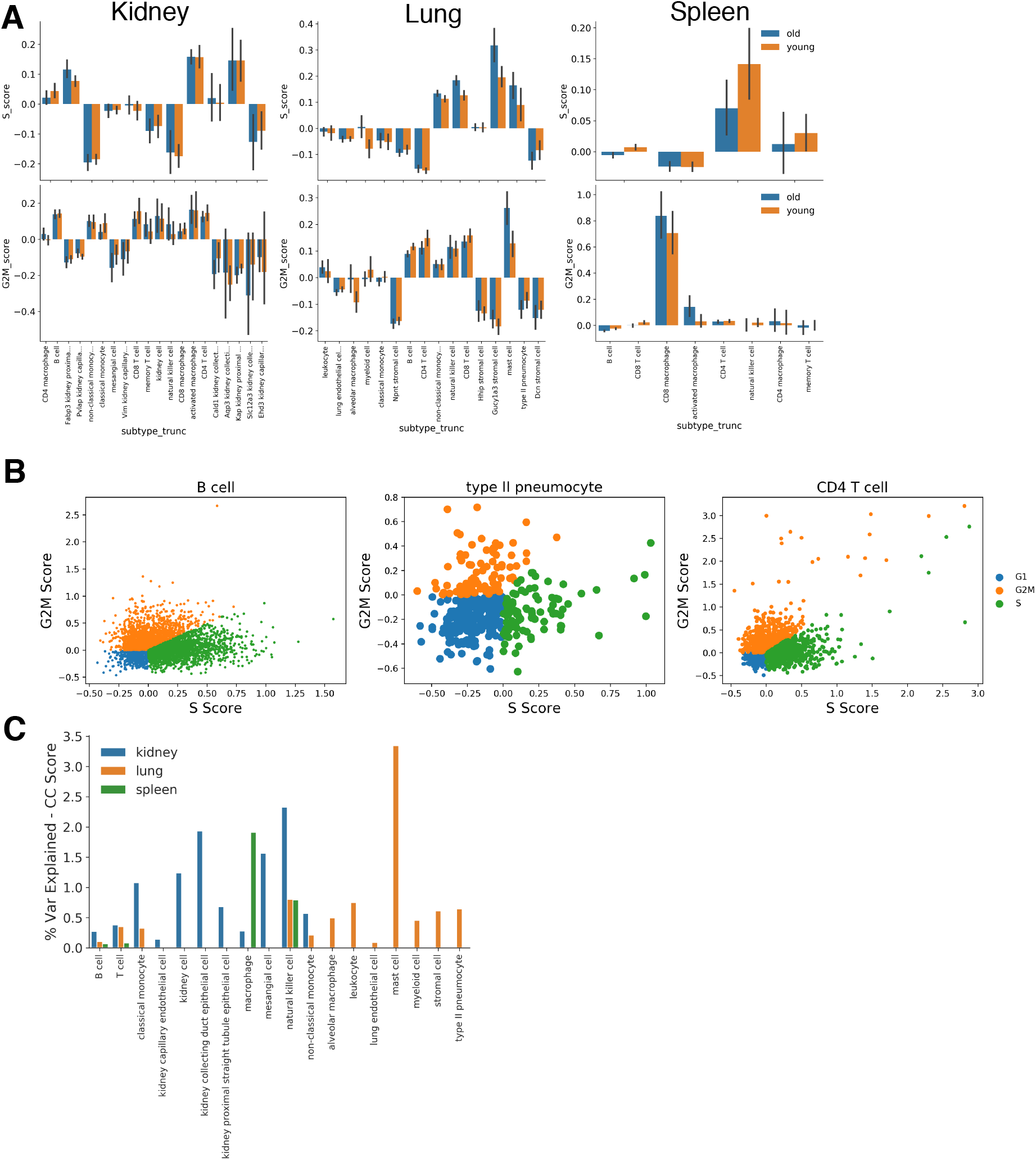
Cell cycle activity is largely unchanged with age. **(A)** S-phase gene module and G2/M-phase gene module scores each cell type in each tissue. We find little difference in CC module scores across cell types. **(B)** Representative cell cycle phase plots based on gene module scoring for S-phage and G2M-phase genes. Here we show individual cell states from the lung. We find very few distinctly cycling cells across all those observed. **(C)** Proportion of variance explained by cell cycle module scores based on linear models.

**Figure S8:**
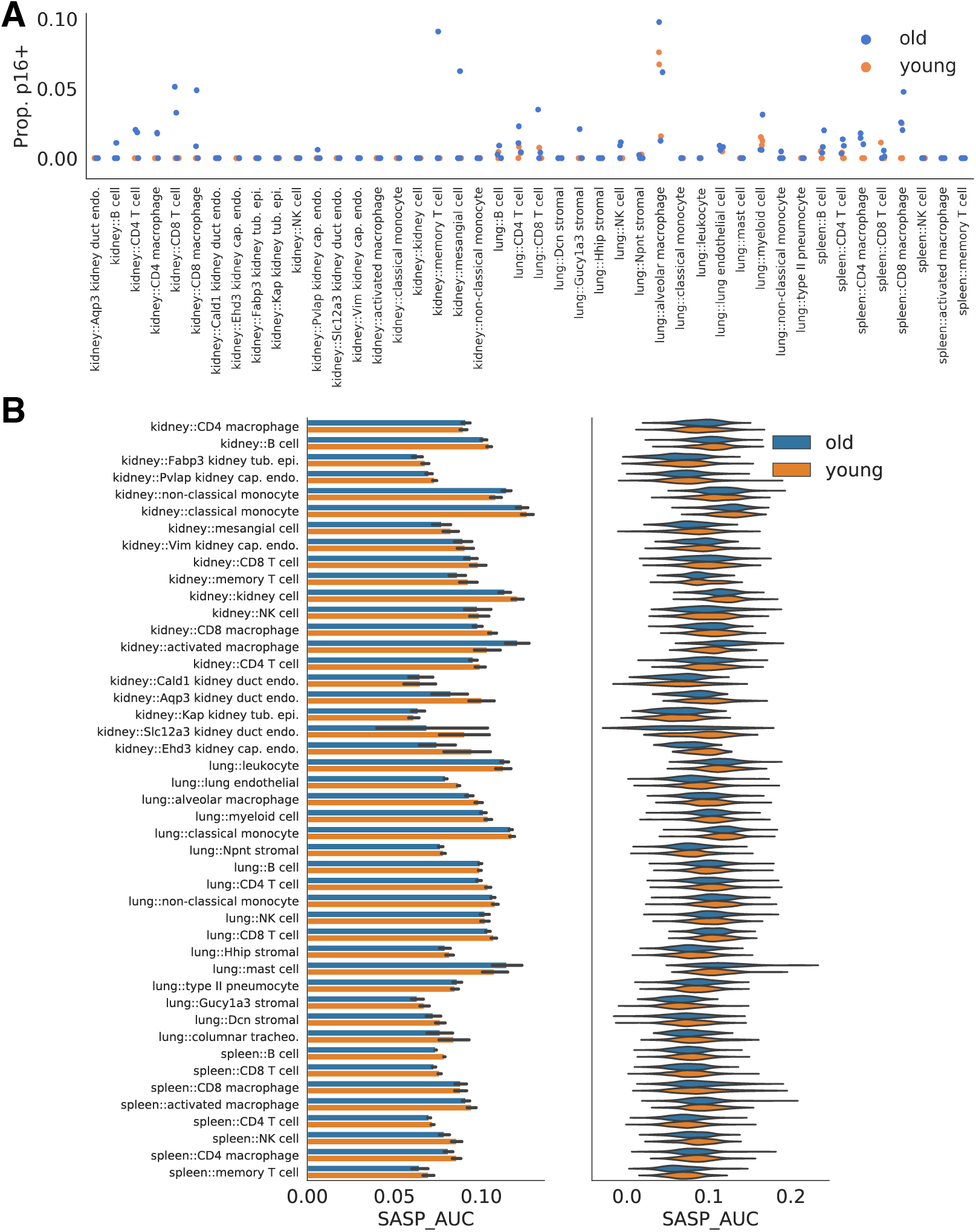
Neither *Cdkn2a* (p16-Ink4a) nor senescence-associated gene activity is significantly upregulated in old cells. **(A)** We quantify the percentage of cells in each identity that express *Cdkn2a* for each animal. No cell identity shows a significantly increased proportion of *Cdkn2a*+ cells. **(B)** We scored the activity of a manually curated set of senescence-associated secretory phenotype (SASP) genes using the AUCell approach for each cell identity at each each. Higher AUCell score indicate high activity of the gene program. We do not find higher activity of the SASP genes in old cells.

**Figure S9:**
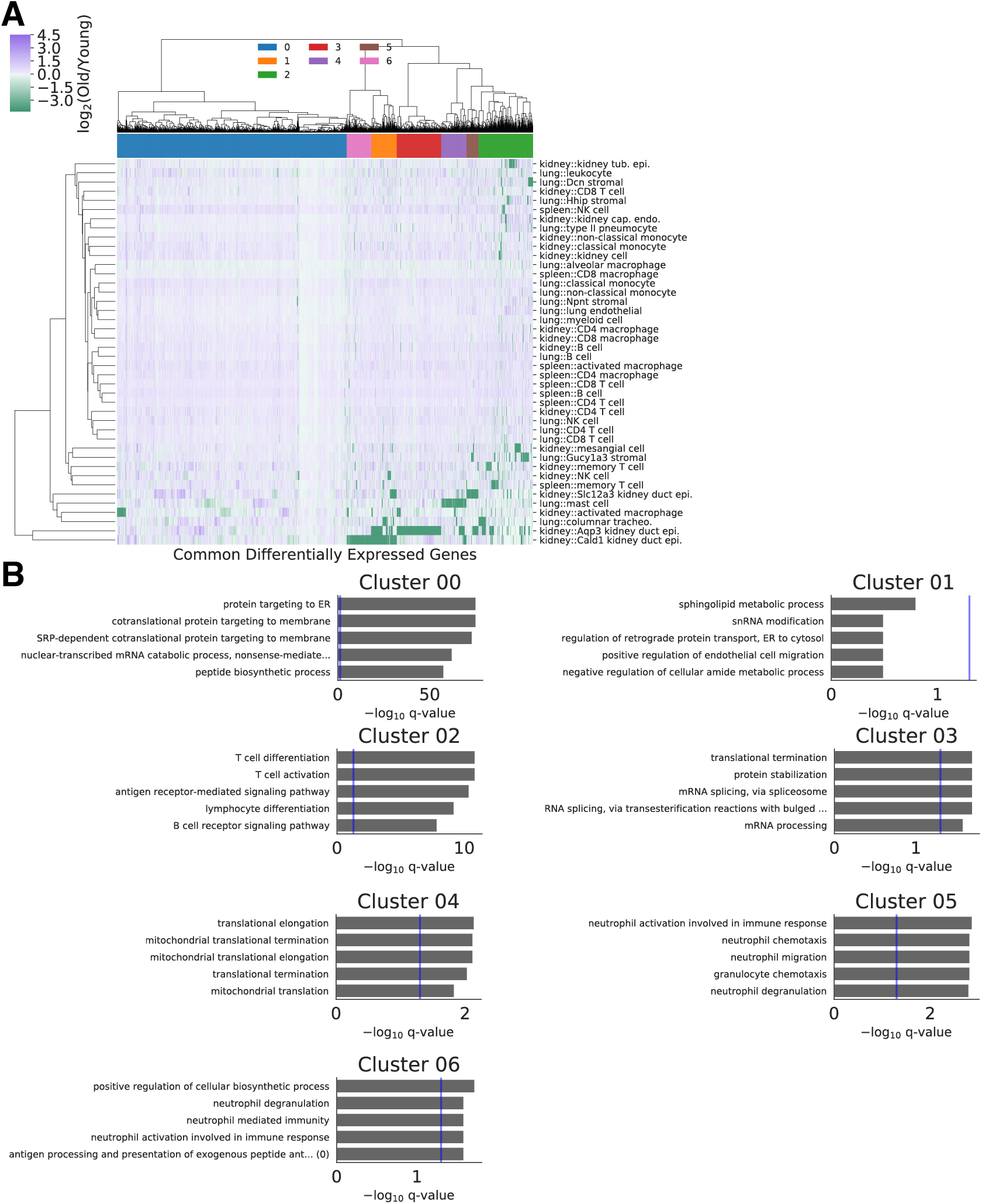
Differential expression identifies a set of genes changed with age across many cell identities. **(A)** We identify a set of 275 genes change with age in at least *k* = 5 cell identities. Performing hierarchical clustering on the fold-changes within each cell identity, we identify subsets of these genes with similar behavior (color column labels). We used cosine similarity as an affinity metric for clustering. **(B)** Gene ontology enrichment terms for genes within each gene cluster identified above.

**Figure S10:**
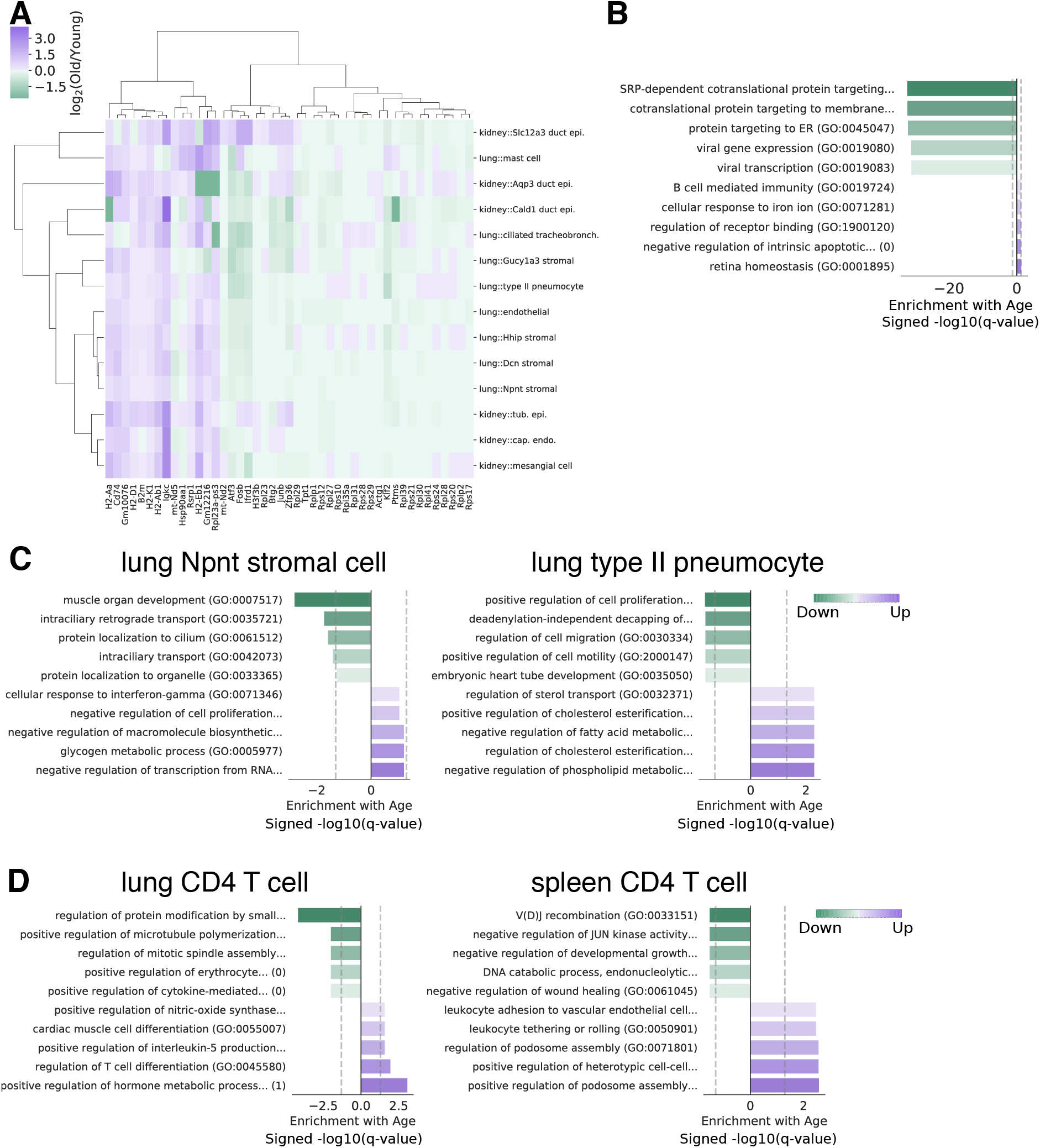
Differential expression analysis identifies age-related changes unique to cell identity and tissue environment. **(A)** Heatmap of common differentially expressed genes found in *k* > 3 non-immune cell states. We note that *B2m, Ikgc*, and *Cd74* are commonly upregulated with aging, even in these non-immune cells. **(B)** Gene ontology enrichment terms for common differentially expressed genes in non-immune cell states. Dashed grey lines demarcate the *α* = 0.05 significance threshold for enrichment. We find that immunological activation pathways are upregulated even in these non-immune cell states, though we note that the gene ontology enrichments are modest. SRP-dependent protein localization and ER targeting are again downregulated. **(C)** Gene ontology enrichment analysis for genes up- and downregulated with age in type II pneumocytes, but not *Npnt* stromal cells and vice-versa. Dashed grey lines demarcate the *α* = 0.05 significance threshold for enrichment. Genes downregulated with age reflect the mesenchymal nature of lung stroma and unique fluid shear stresses in type II pneumocytes. **(D)** Gene ontology enrichment analysis for genes up- and downregulated with age in natural killer cells from the lung, but not natural killer cells from the spleen and vice-versa.

**Figure S11:**
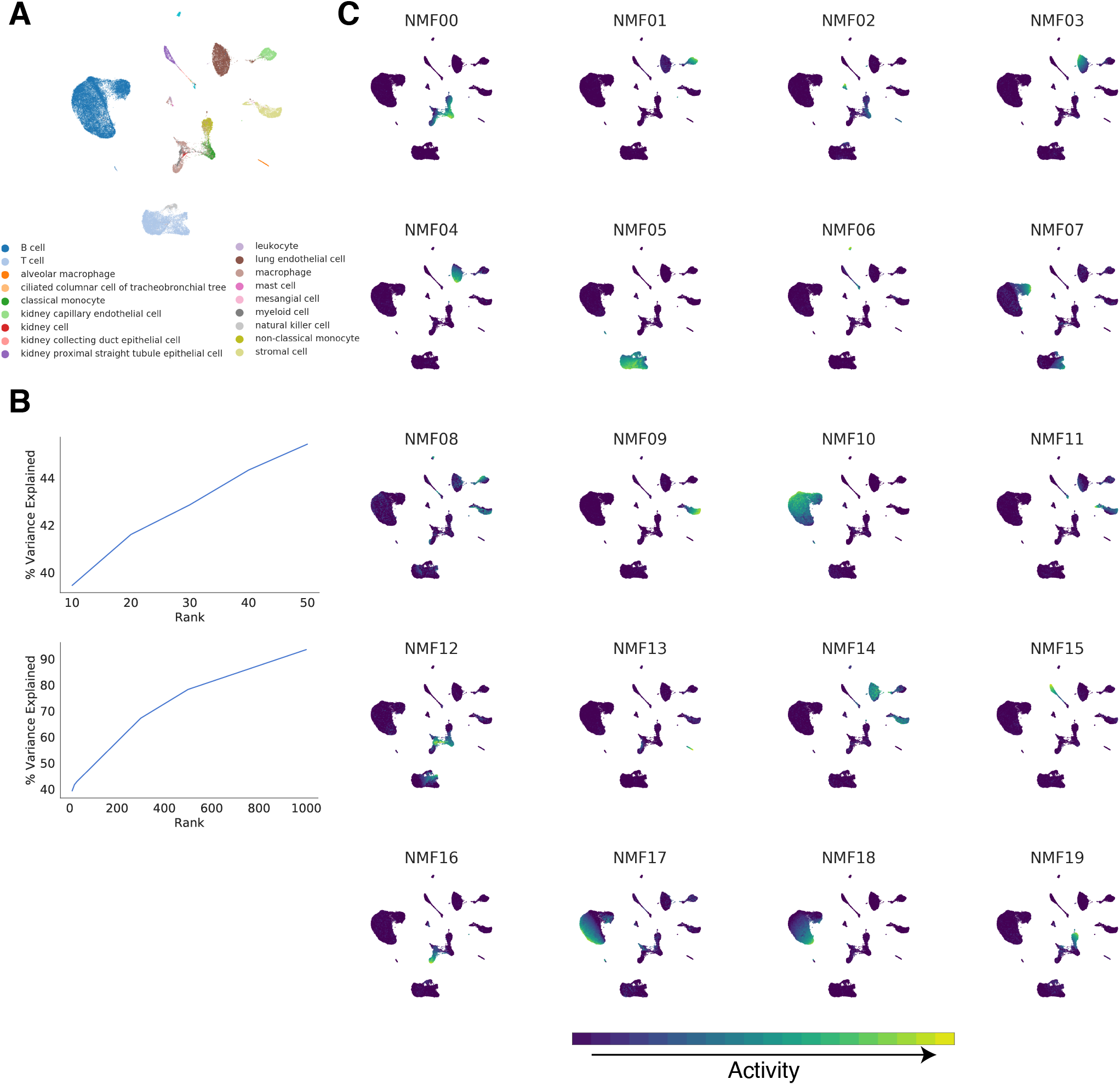
Non-negative matrix factorization of all observed cell types. **(A)** UMAP projection of our NMF embedding with rank *k* = 20. **(B)** Proportion of variation in count data explained by NMF embeddings as a function of rank and initialization procedure. We find that a rank of *k* = 20 sits at the elbow of this relationship, capturing ≈ 42% of the variation. **(C)** We visualize the activity level of each gene expression program captured by the NMF embedding across cell types in a UMAP projection. Note that the NMF dimensions are not explicitly ordered.

**Figure S12:**
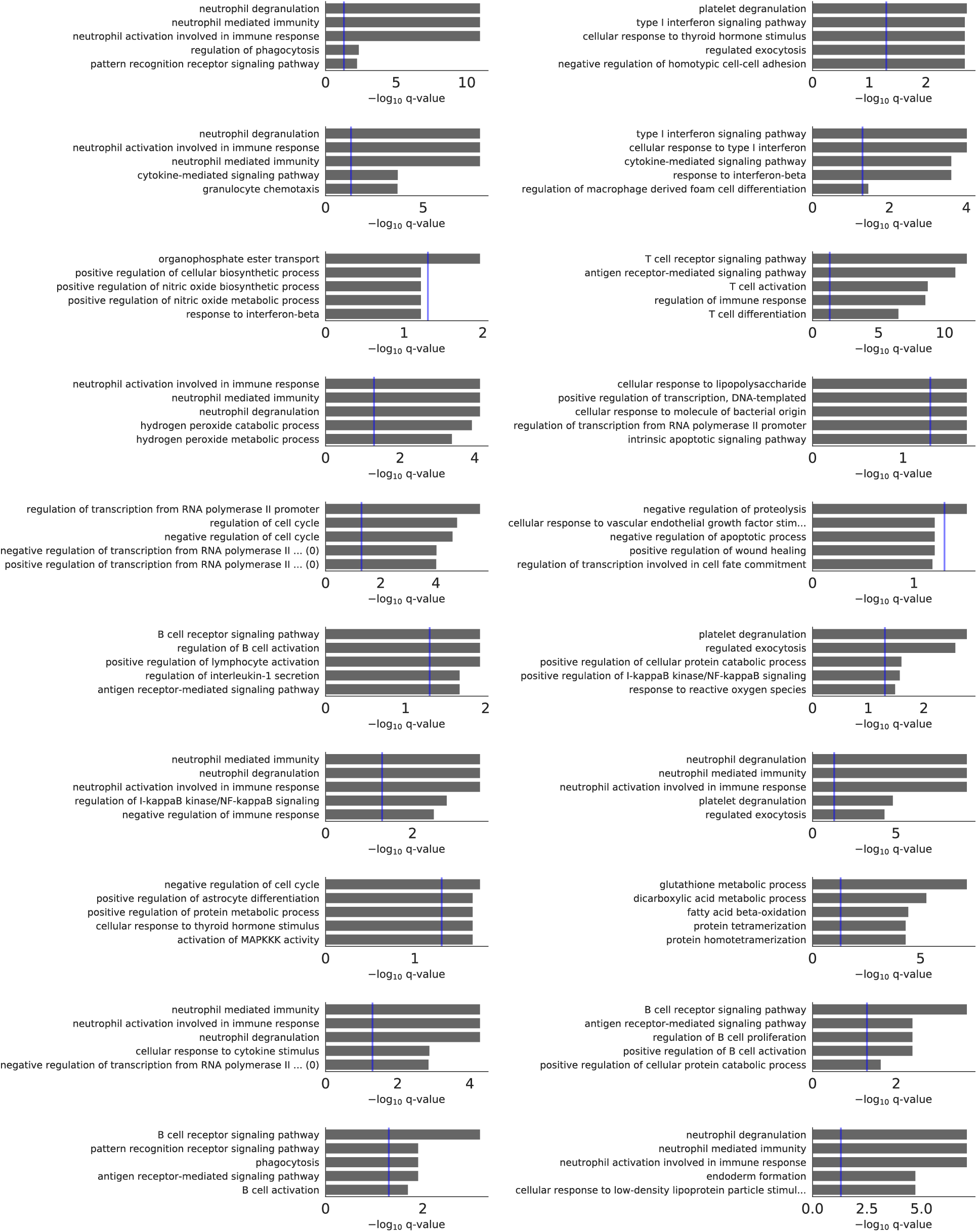
Gene ontology enrichment in non-negative matrix factorization embedding dimensions. Top 10 enriched Gene Ontology Terms for each dimension of the NMF embedding. We performed Gene Ontology enrichment analysis on the genes with loadings above a threshold value for each dimension.

**Figure S13:**
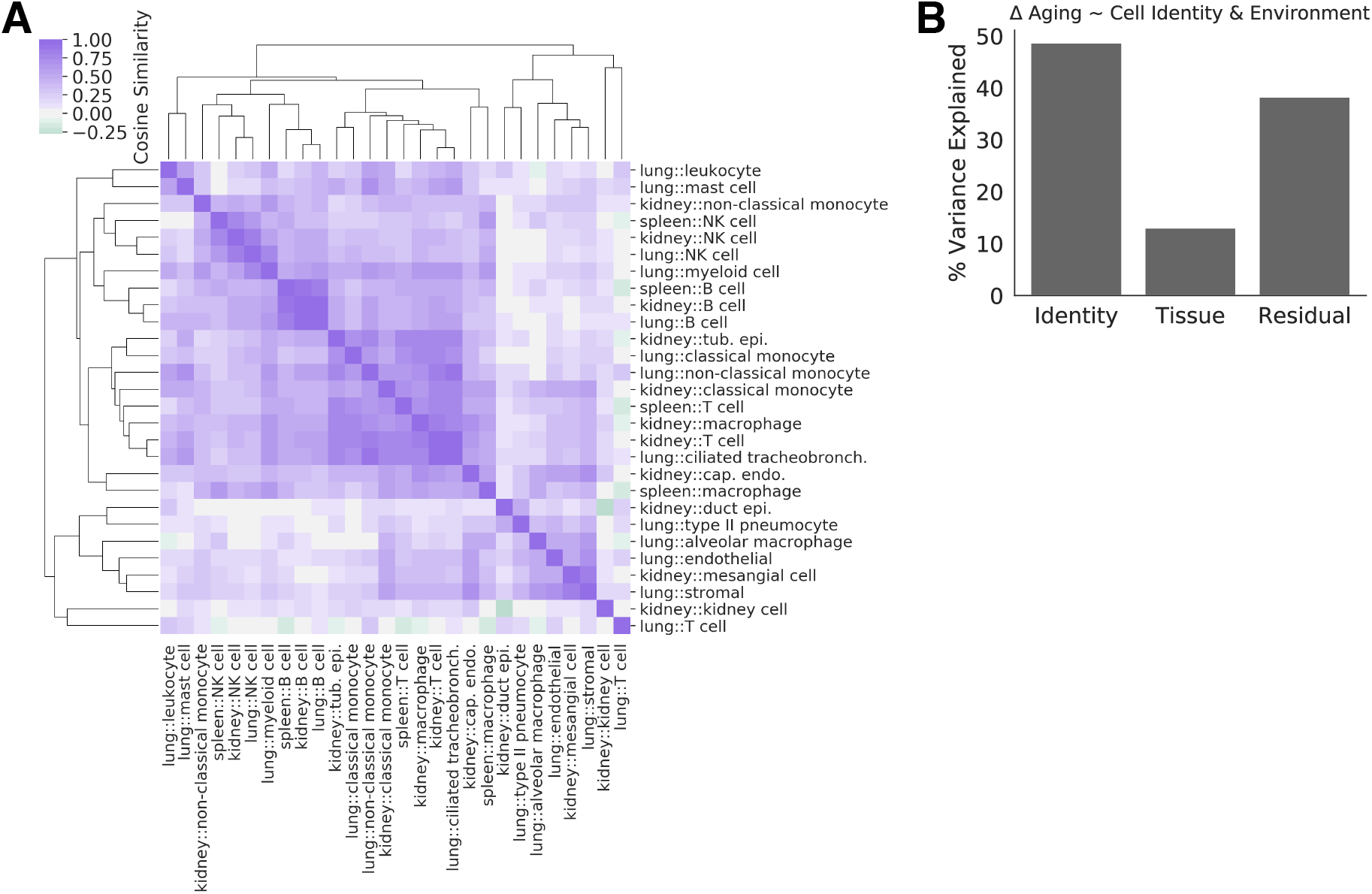
Cell identity and tissue environment influence on aging trajectories in a PCA embedding. **(A)** We computed aging trajectories between young and aged cell centroids in a PCA embedding. We compare these trajectories using cosine similarities. Cosine similarities between the aging trajectories of each cell state in each tissue are presented as values in the heatmap. **(C)** Variation in the aging vectors (computed in the PCA embedding) of immune cell types found in all three tissues explained by cell type and tissue environment (ANOVA).

**Figure S14:**
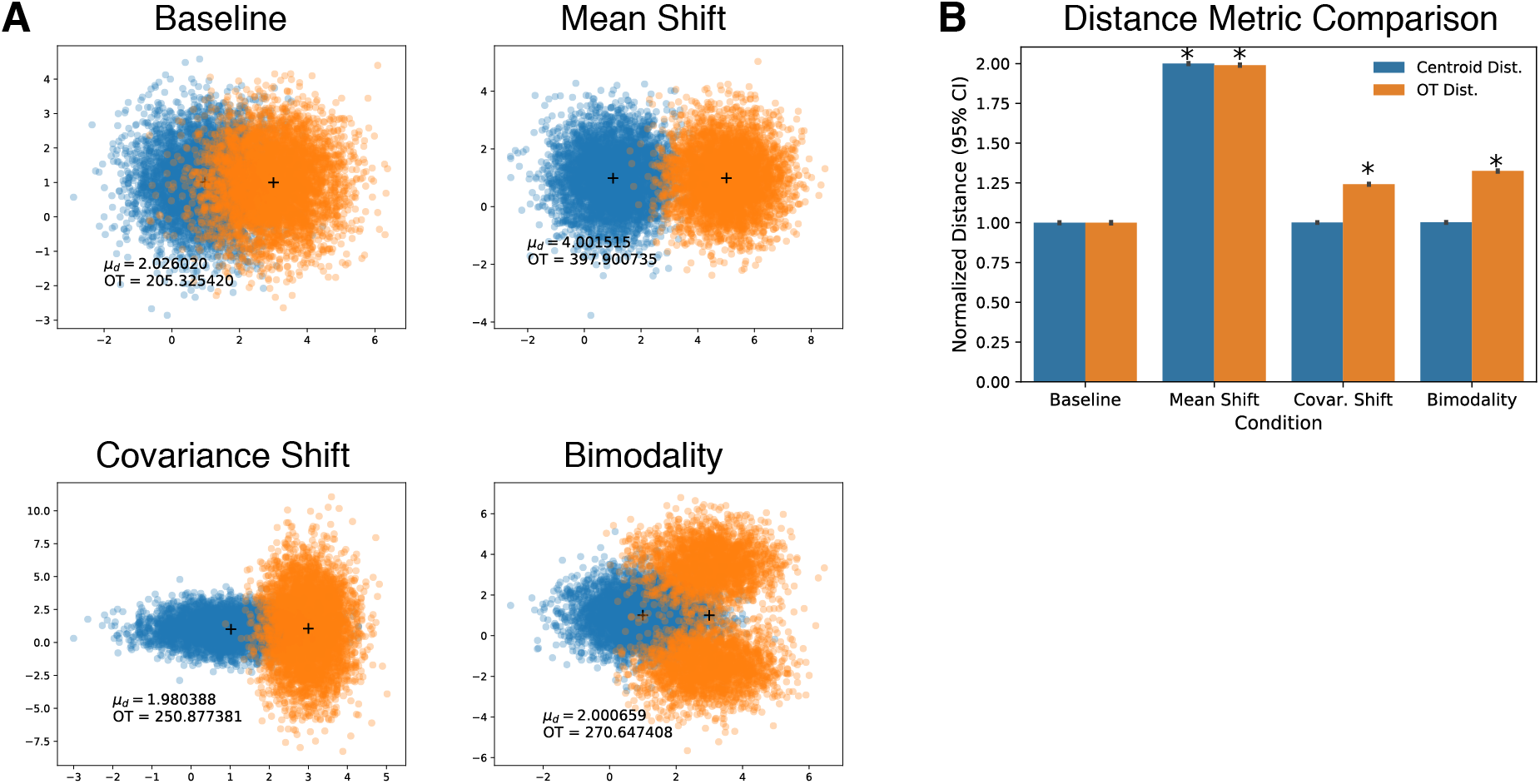
Simulation experiments demonstrate that optimal transport distances capture differences between cell populations. **(A)** Two “cell populations” (blue, orange) were simulated as 2-dimensional Gaussian distributions. We computed the centroid distance and optimal transport (OT) distance between populations for a range of possible differences that may arise between cell populations (text insets). As a baseline, we simulate unit Gaussians with different means. When the difference in means is increased (Mean Shift), both the centroid distance and OT distance reflect the magnitude of change. However, when we shift the covariance matrix of one population or simulate a bimodal population with the same mean as the baseline unimodal population, only the OT distance reflects these differences. **(B)** Comparison of centroid distance and OT distance metrics for comparing simulated cell populations. Each simulated population contains *n* = 5000 cells. Each simulation was performed 50 times to estimate confidence intervals. For each iteration of the simulation, we compute the OT distance as the mean OT distance across 30 random samples of *n* = 100 cells from each simulated population. (*: *p*-value < 0.05, *t*-test to baseline).

**Figure S15:**
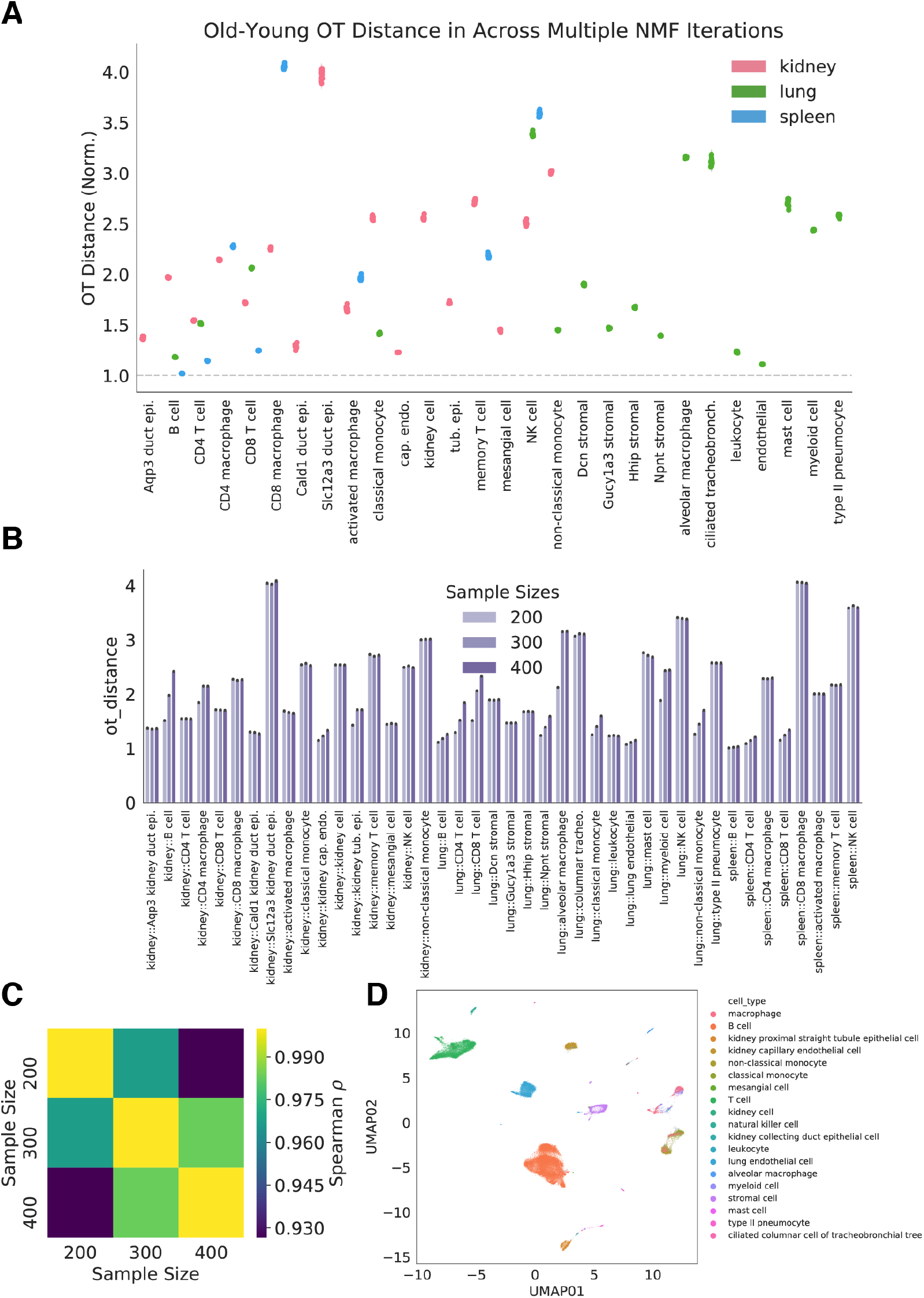
Optimal transport estimates the magnitude of aging across cell identities. **(A)** Optimal transport distances were computed for each cell identity tissue combination. We normalize our heterochronic comparison (Young-Old cells) by the larger mean of two null isochronic comparisons (Young-Young, Old-Old). We computed this normalized distance for 10 separate NMF optimizations. The mean normalized distance for a single NMF optimization is represented as a point, with violins outlining the distribution across 10 iterations. We find that the relative distances between cell identity/tissue combinations are not changed across NMF optimization runs. **(B)** Optimal transport distances were computed for a range of random sampling sizes. We present the normalized Old-Young distance values (normalized as in **(A)**) for each sample size. **(C)** Heatmap of Spearman correlations between the normalized Old-Young distances computed using different sample sizes. Old-Young distances have high correlation (Spearman’s *ρ* > 0.9) across the range of sample sizes, suggesting the metric is robust to changes in sample size. **(D)** UMAP projection of the NMF (rank 500) embedding used for optimal transport distance calculation. Cell types are overlaid as colors.

### Supplemental Tables

**Table S1:**
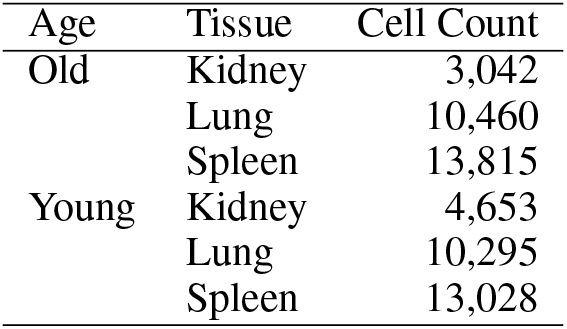
Total cell counts for each tissue, stratified by age.

## Notes

http://mca.research.calicolabs.com

